# Distinct effects of acute and chronic blood loss anemia on vascular function after acute myocardial infarction

**DOI:** 10.1101/2024.09.24.614629

**Authors:** Isabella Solga, Aslihan Sahin, Vithya Yogathasan, Lina Hofer, Feyza Gül Celik, Amira El Rai, Mohammed Rabiul Hosen, Patricia Wischmann, Stefanie Becher, Amin Polzin, Norbert Gerdes, Christian Jung, Malte Kelm, Ramesh Chennupati

## Abstract

**Background:** Anemia is frequently observed in patients with cardiovascular diseases (CVD). Anemia alone or in combination with other morbid conditions leads to poor prognosis in acute myocardial infarction (AMI). We recently showed that moderate blood loss anemia is associated with red blood cell (RBC) dysfunction and a compensatory increase in flow-mediated dilation (FMD) responses which are compromised in chronic blood loss anemia However, the effects of acute anemia (AA) and chronic anemia (CA) on endothelial function after AMI are unclear. In this study, we evaluated systemic endothelial function following AMI in established murine models of blood loss acute and chronic anemia. We hypothesize that both AA and CA aggravate systemic endothelial dysfunction (ED) after AMI.

**Methods and results:** AA or CA was induced in male C57BL/6J mice by repeated blood withdrawal for three consecutive days or six weeks, respectively. Separate groups of anemic and non-anemic mice underwent AMI via left anterior descending artery (LAD) ligation (45 min), followed by reperfusion. Endothelial function was assessed using both *in vivo* and *in vitro* methods 24 h post-AMI. Impaired flow-mediated dilation (FMD, *in vivo*) and endothelium-dependent relaxation (EDR) responses were observed in the aorta, femoral, and saphenous arteries of AA mice compared to their respective control groups 24 h post AMI. The aorta and saphenous arteries from CA mice showed significantly reduced vascular smooth muscle (VSM) contractile responses after AMI. Analysis of oxidative products of nitric oxide (NO) in plasma revealed reduced nitrite and nitrate levels in both AA and CA mice compared to controls 24 h post-AMI. Immunohistochemistry of aortic tissues from both anemic groups showed increased reactive oxygen species (ROS) product 4-Hydroxynonenal (4-HNE). Co-incubation of RBCs from anemic mice or anemic ST-elevation myocardial infarction (STEMI) patients with aortic rings from wild type mice demonstrated attenuated VSM contractile and EDR responses. Supplementation with the ROS scavenger N-acetyl cysteine (NAC) for four weeks improved both *in vivo* and *ex vivo* EDR in AA and CA mice 24 h post-AMI.

**Conclusion:** After AMI, both AA and CA are associated with severe ED, while VSM contractile responses specifically reduced in CA mice. These effects are accompanied by increased ROS and partly mediated by RBCs. Antioxidant supplementation with NAC is a potential therapeutic option to reverse the severe vascular dysfunction in anemia following AMI.

**Graphical Abstract:** **Distinct effects of acute and chronic anemia on vascular function 24 h post-AMI.** After acute myocardial infarction, acute and chronic anemia are associated with increased reactive oxygen species (ROS) and inflammation in endothelial cells (EC), leading to the inhibition of endothelial nitric oxide synthase (eNOS) and subsequent endothelial dysfunction by limiting NO bioavailability. Chronic anemia is additionally associated with decreased vascular smooth muscle cell (VSMC) function due to increased oxidative stress, leading to SMC dysfunction. After N-Acetyl-L-Cysteine (NAC) treatment, vascular function is improved in both anemic groups.

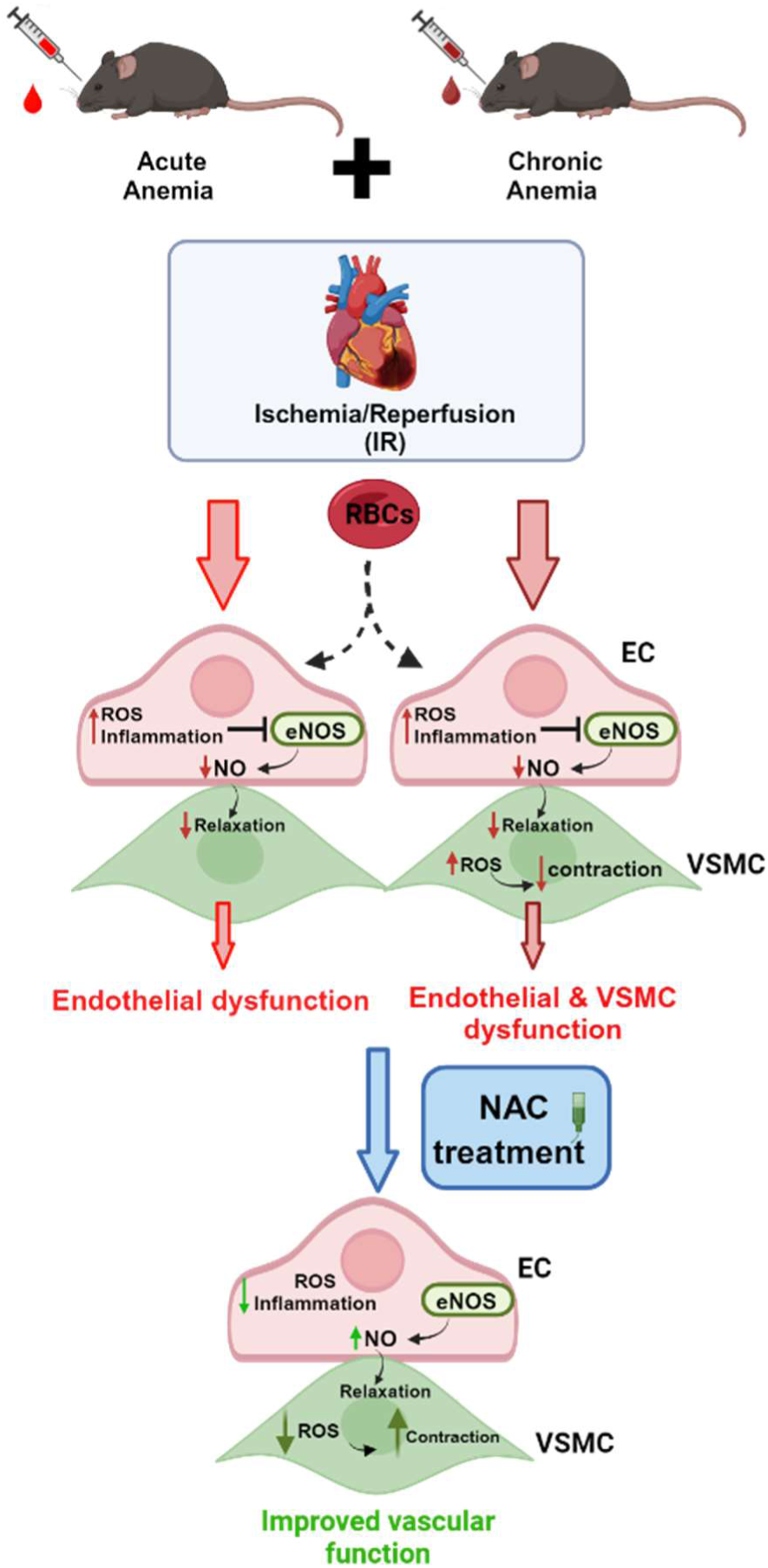

## Introduction

Acute myocardial infarction (AMI) is one of the leading causes of sudden death worldwide due to a blockage in the coronary arteries resulting in necrosis of the myocardium (*1–3*). Anemia is a frequently diagnosed co-morbidity in patients with AMI, characterized by reduced hemoglobin levels, hematocrit, and a reduction in circulating red blood cells (RBCs). Many elderly patients with cardiovascular disease develop anemia during hospitalization due to blood loss during interventions or diagnoses, which is defined as hospital-acquired anemia which might have consequences on the vascular system and thus on prognosis during cardiovascular events (*4*). Anemia in association with acute coronary syndrome, stroke, or heart failure leads to poor cardiovascular disease prognosis (*5*). Furthermore, anemic patients have a higher risk of major bleeding, arrhythmias, and heart failure after AMI, leading to morbidity and mortality (*6, 7*). However, the underlying mechanisms of how hospital-acquired anemia influences vascular function is largely unknown.

The endothelium in arteries plays a fundamental role in regulating vascular tone by synthesizing and releasing an array of endothelium-derived relaxing factors, such as nitric oxide (NO) and endothelium-derived hyperpolarizing factor (EDHF). NO is generated from L-arginine by endothelial NO synthase (eNOS), which diffuses into vascular smooth muscle cells (VSMCs) and activates guanylate cyclase, resulting in cyclic guanosine monophosphate (cGMP)-mediated vasodilation (*8*). Reduced production of endothelium-dependent NO and increased levels of reactive oxygen species (ROS) and inflammation are associated with many forms of cardiovascular diseases, including hypertension, coronary artery disease, and chronic heart failure, due to increased oxidative stress (*9–11*). It is well known that increased ROS in the endothelium uncouple eNOS, leading to reduced NO production and thus endothelial dysfunction (ED). ED is a hallmark of vascular dysfunction in AMI. Clinically, the assessment of flow-mediated dilation (FMD) is the gold standard for evaluating endothelial function *in vivo*, which is primarily mediated by NO (*12*). Decreased FMD responses are associated with various cardiometabolic diseases, such as myocardial infarction, diabetes, and coronary artery disease (*13*). Previous studies have shown that anemia is associated with increased FMD responses in humans without comorbidities (*14*). Anemia-related hemoglobinopathies, or anemia in combination with diabetes or chronic kidney disease (CKD), are associated with reduced FMD responses (*15–17*). These studies demonstrate that endothelial function is altered based on the type of anemia and associated co-morbid conditions. Additionally, we recently showed that acute blood loss anemia is associated with increased compensatory FMD responses, which are compromised in cases of chronic anemia (CA) due to increased oxidative stress (*18, 19*). In the same study, we demonstrated that CA is associated with progressive ED in large arteries due to increased oxidative stress in the endothelium (*19*). However, the effect of anemia on endothelial function after AMI remains largely unknown.

RBCs are known to induce ED in different disease states such as diabetes, hypocholesteremia, preeclampsia, and anemia (*19–22*). We have recently shown that RBCs in anemia lose their cardioprotective properties in ST-elevation myocardial infarction (STEMI) patients (*23*), demonstrating the potential role of RBCs in cardiometabolic diseases. It is not clear how acute and chronic blood loss anemia affects vascular function after myocardial infarction, and the potential role of RBCs in this has never been investigated. Therefore, in this study, we used two well-established acute and chronic blood-loss mouse models to study the effect of blood loss anemia on vascular function, namely, endothelial and VSMCs after Ischemia-Reperfusion (IR) injury. Additionally, we investigated whether RBCs from anemic mice with AMI and anemic STEMI patients affect endothelial function using co-incubation with aortic rings followed by VSCM and endothelial function analysis. To our knowledge, this is the first study to evaluate systemic ED after AMI in acute and chronic blood-loss anemia murine models.

## Materials and Methods

### Animals

All animal procedures used in the study were approved and performed in accordance with the ARRIVE (Animal Research: Reporting of *In Vivo* Experiments) II guidelines and authorized by LANUV (North Rhine-Westphalia State Agency for Nature, Environment and Consumer Protection) in compliance with the European Convention for the Protection of Vertebrate Animals used for Experimental and other Scientific Purposes. The approval numbers for the animal experiments are 84-02.04.2020.A073 and 84-02.04.2018.A234. The C57Bl/6J (wildtype, WT) mice were obtained from Janvier Labs (Saint-Berthevin Cedex, France). Mice were housed in standard cages (constant room temperature and humidity, with 12h light/dark cycles) and had free access to standard pelleted food and tap water.

### Anemia induction

For the experimental approach, we used four different groups of mice. The mice were divided into AA, CA, and their respective non-anemic groups. AA was induced in 10- to 12-week-old male C57BL/6J wildtype (WT) mice by repetitive mild blood withdrawal on three consecutive days, resulting in <20 g/l changes in hemoglobin (Hb). The blood was withdrawn from the facial vein under isoflurane anesthesia (3%). The amount of daily blood loss per mouse was adjusted to <15% of the total blood volume and was replaced by saline administration. CA was induced by mild blood withdrawal every third day for a period of six weeks, also resulting in <20 g/l changes in Hb. At the end of the sixth week, mice with hemoglobin levels <10 g/l were considered for experiments. The two non-anemic groups were age-matched to the anemic groups and underwent the same handling as the anemic mice (including puncture of the facial venin) without blood withdrawal.

### Induction of AMI

In a separate set of anemic and non-anemic groups of mice, acute myocardial infarction (AMI) was induced. Briefly, mice were anesthetized with isoflurane, intubated, and ventilated with a tidal volume of 0.2-0.25 mL and a respiratory rate of 140 breaths per minute, using isoflurane (3%) and 30% O_2_ with a rodent ventilator. Body temperature was controlled and maintained at 37°C throughout the surgical procedure. A left lateral thoracotomy was performed between the third and fourth rib, and the pericardium was exposed and dissected. Ischemia was induced by gently tightening a 7-0 surgical suture placed under the left anterior descending (LAD) artery. To ensure that the procedure was carried out correctly, in addition to the visible blanching of the apex, changes in the electrocardiogram (ECG; ST-segment elevation) were monitored. After 45 minutes, the ligation was removed, and the myocardium was reperfused for 24 h. Animals received buprenorphine (0.5 mg/kg BW) subcutaneously every 8 h until euthanasia.

### Supplementation of NAC

To investigate the potential role of reactive oxygen species (ROS) in mediating endothelial dysfunction, additional groups of mice were supplemented with 1% N-acetylcysteine (NAC) (Sigma) through drinking water for 4 weeks. After 4 weeks, acute or chronic anemia was induced, and ischemia-reperfusion (IR) surgery was performed to induce AMI.

### Collection of Blood from STEMI Patients

Blood was collected from STEMI patients, with and without anemia, within 24 h post-infarction in EDTA tubes. The patients were diagnosed based on changes in the ECG. According to the World Health Organization (WHO) guidelines, adult male patients with hemoglobin (Hb) levels below 13.0 g/dL and adult female patients with Hb levels below 12.0 g/dL are considered anemic (*24*). All patients included in this study provided written consent and were recruited from the Department of Cardiology, Pulmonology, and Angiology at Düsseldorf University. Permission numbers for blood sample collections are 5481R, 2018-14, and 2018-47, as approved by the ethics committee of Düsseldorf University Hospital.

### *In vitro* studies with isolated aortic rings

#### Solutions and drugs

Krebs-Ringer bicarbonate-buffered salt solution (KRB) contained (in mmol/L): 118.5 NaCl, 4.7 KCl, 2.5 CaCl_2_, 1.2 MgSO_4_, 1.2 KH_2_PO_4_, 25.0 NaHCO_3_ and 5.5 glucose. The KRB solution was continuously aerated with 95% O_2_/5% CO_2_ and maintained at 37°C. Indomethacin (INDO; Sigma Aldrich,) was dissolved in ethanol. Acetylcholine (ACh), phenylephrine (PHE), N**^ω^**-nitro-arginine methyl ester (L-NAME) and sodium nitroprusside (SNP; all Sigma Aldrich) were dissolved in KRB solution.

#### Organ chamber experiments

Mice were euthanized under deep isoflurane anesthesia (4.5%). The thoracic aorta was dissected free from perivascular adipose tissue and cut into 2 mm aortic rings. The segments were then mounted in a wire myograph system (Danish Myo Technology (DMT), Aarhus, Denmark) and stretched to a force of 9.8 mN. The segments were allowed to normalize for 45 minutes with a periodic buffer change three times. Saphenous and femoral arteries were dissected free from surrounding fat and connective tissue and mounted in a wire myograph (DMT). Arterial segments (2 mm) were stretched to the optimal diameter at which maximal contractile responses to 10 μM norepinephrine (NA) could be obtained (*25*).

##### Contributions of NO, cyclo­oxygenase products to endothelium­dependent relaxation

In the first step, a concentration-response curve (CRC) for PHE (0.001-10 μM) and ACh (0.001-10 μM) was generated in presence of the cyclooxygenase inhibitor indomethacin (INDO, 10 µM). Next, to evaluate the contribution of NO to the relaxation response in the arteries the CRC was repeated in the presence of both INDO and L-NAME (100 µM) a NOS inhibitor.

##### Sensitivity of vascular smooth muscle to NO

Additionally, the SMC sensitivity to NO was evaluated by performing a CRC in presence of INDO and L-NAME (100 µM), with the NO donor SNP (0.01-10 µM).

#### Co-incubation experiments with murine and human RBCs

For further investigations of dysfunctional RBCs on endothelial-dependent relaxation responses in anemia, we performed co-incubation studies with murine and human RBCs. Blood was collected in EDTA tubes from non-anemic and anemic mice 24 h post-AMI, and additionally from STEMI patients with and without anemia. The RBCs were isolated by centrifugation at 800 xg for 10 minutes at 4°C, and 40% hematocrit (hct) was prepared using KRB buffer. Aortic rings (2 mm) isolated from WT mice were co-incubated with the isolated RBCs for 2 h (murine RBCs) or 6 h (human RBCs) at 37°C. After the incubation, the aortic segments were mounted in the wire myograph system, and vascular function was evaluated.

### Flow-mediated dilation assessment

The C57BL/6J mice of week-old mice with and without anemia were used for the assessment of flow-mediated dilation (FMD) responses. The measurements were performed 24 h post - AMI. FMD responses were determined by using a Vevo 2100 high-resolution ultrasound scanner using a 30–70 MHz linear transducer (Visual Sonics Inc., Toronto, Canada) as described previously (*26*). During the whole procedure/measurement mice were kept under 2-2.5% isoflurane anaesthesia and a body temperature of 37°C was maintained. To visualise the femoral artery, the transducer was placed at the lower limb and a vascular occluder (8 mm diameter, Harvard Apparatus, Harvard, Boston, MA, USA) was used to perform the dilation measurement (*26*). Initially baseline images of the vessel were recorded. In order to measure changes in the vessel diameter, first an occlusion phase was performed by inflating the cuff to 250 mmHg. During the occlusion the pressure was kept constant for 5 min and an image was taken every 30 s (Druckkalibriergerät KAL 84, Halstrup Walcher, Kirchzarten, Germany). Afterwards the cuff was released (reperfusion phase) for 5 min and again every 30 s an image was taken to determinate the FMD. Changes in vessel diameter were quantified as percent of baseline (%) = [diameter (max)/diameter (baseline)] × 100.

### Immunohistochemistry

Thoracic aortas from the anemic mice 24 h post AMI and respective non-anemic mice 24 h post AMI were fixed in formaldehyde (4%) for 2 h and stored in sucrose (30%) overnight. Afterwards the tissues were embedded in Tissue-Tek®, O.C.T and frozen until further use. Tissue sections (5 µm) were incubated overnight at 4 °C with rat anti-mouse CD31 (1:200 in blocking solution ((0.1% Saponin, 0.5% BSA, 0.2% fish gelatin in 1 x PBS) BD Biosciences) and goat anti-rabbit 4-HNE (1:200 in blocking solution; Abcam) antibodies. Next, the sections were incubated with respective goat anti-rat Alexa Fluor 488 (1:1000 in blocking solution; ThermoFisher) and goat anti-rabbit Alexa Fluor 555 (1:1000 in blocking solution; ThermoFisher) secondary antibodies. Additionally, autofluorescnece was quenched using Vector® TrueVIEW® Autofluorescence Quenching Kit (BIOZOL Vectorlabs). Prior to covering, a DAPI staining (1mg/ml) was performed for 5 min and then the sections were mounted with VectaMount® Aqueous Mounting Medium (ThermoFisher) and imaged with Leica DM6 M microscope (Leica Microsystems).

### Assessment of plasma inflammatory markers

Blood was collected from all experimental groups in EDTA tubes. RBC and plasma samples were prepared and snap-frozen. The samples were stored at −80°C until further use. Plasma samples were used to assess inflammatory parameters in the different mice groups by using ELISA kits for VCAM-1 and ICAM-1 measurements (R&D Systems).

## Results

### Acute anemia is associated with altered endothelial function after AMI

To assess the effect of AA on vascular function 24 h post-AMI, contractile responses, and endothelium-dependent and -independent relaxation responses were measured in isolated aortic rings (large artery) using wire myograph. The contractile responses to phenylephrine did not significantly differ between AA mice and non-anemic mice 24 h post-AMI (Suppl. Table 1). However, the endothelium-dependent relaxation responses to acetylcholine in the presence of indomethacin were significantly reduced in AA mice (E_max_: 25.22 ± 2.88%; p=0.0009) compared to the corresponding non-anemic group (E_max_: 52.45 ± 6.02%) of mice 24 h post-AMI (Fig. 1A; Suppl. Fig. 1A; Suppl. Table 1). In the presence of the NOS inhibitor (L-NAME, 100 µM), relaxation responses were completely inhibited in both AA and non-anemic groups of mice (Fig. 1B), proving that the relaxation responses are mediated by NO. SMC sensitivity to NO and endothelium-independent relaxation were assessed using an exogenous NO donor, SNP. All groups showed similar relaxation responses, indicating that SMC sensitivity to NO was similar in both groups (Fig. 1C; Suppl. Table 1).

**Figure 1:**
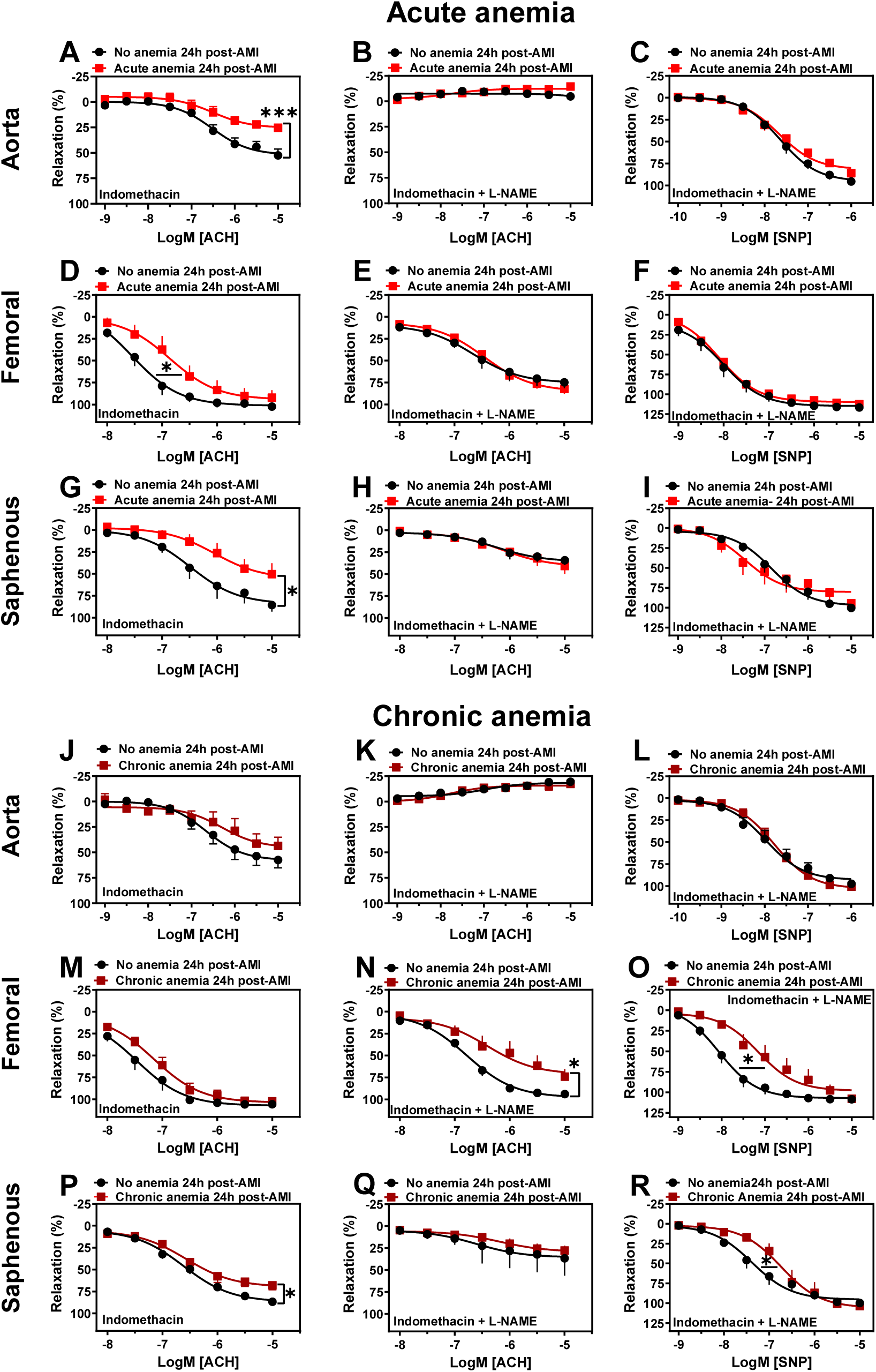
Both acute and chronic anemia are associated with vascular dysfunction 24 h post-AMI. Arterial segments of aorta **(A-C, J-L)**, femoral **(D-F, M-O)** and saphenous arteries **(G-I, P-R)** were isolated from acute (red squares) and chronic (dark red squares) anemic mice and respective non-anemic mice (black circles) 24 h post IR. Arterial segments were pre-contracted using phenylephrine (PHE; 10 µM), and their relaxation responses to acetylcholine (ACh; 1 nM-10 µM) were assessed using wire myography. **(A, D, G, J, M, P)** Relaxation (%) in the presence of indomethacin (10 µM, a COX inhibitor). **(B, E, H, K, N, Q)** Relaxation in the presence of indomethacin and L-NAME (100 µM, NOS inhibitor). Maximal endothelial-dependent relaxation response (E_max, %_). **(C, F, I, L, O, R)** Relaxation responses to sodium nitroprusside (SNP, 10 nM-10 µM) in the presence of indomethacin and L-NAME. All values are mean values ± SEM (n = 8–10 per group). *, p ≤ 0.05; **, p ≤ 0.01; ***, p ≤ 0.001. Concentration-response curves (CRCs) were analysed by Two-Way ANOVA and Bonferroni ‘s post-hoc test to compare acute, chronic and respective non-anemic group of mice.

Vascular function was also assessed in femoral (medium sized) and saphenous arteries (small sized) to investigate possible effects of anemia on different vascular beds. In both femoral and saphenous arteries, the contractile responses to phenylephrine did not significantly differ between AA mice and their respective control group 24 h post-AMI (Suppl. Table 2-3). In the presence of indomethacin, femoral arteries showed reduced sensitivity to acetylcholine-mediated relaxation responses in AA mice (pEC_50_: 6.86 ± 0.24; p=0.04) compared to non-anemic mice (pEC_50_: 7.55 ± 0.20) 24 h post-AMI (Fig. 1D; Suppl. Fig. 1B; Suppl. Table 2). In the small resistance saphenous artery, the endothelium-dependent relaxation responses to acetylcholine were significantly reduced in AA mice (E_max_: 50.36 ± 12.41%; p=0.03) compared to the non-anemic group of mice (E_max_: 85.44 ± 7.59%) 24 h post-AMI (Fig. 1G; Suppl. Fig. 1C; Suppl. Table 3). In both femoral and saphenous arteries, in the presence of L-NAME, the relaxation responses were similar between anemic and non-anemic groups (Fig. 1E & H; Suppl. Tables 2-3). Similarly, endothelium-independent relaxation responses to SNP did not differ between AA mice and their respective control groups 24 h post-AMI (Fig. 1F & I; Suppl. Tables 2-3). These results conclude that AA is associated with reduced endothelium-dependent relaxation responses, which are mainly mediated by NO, despite the smooth muscle sensitivity to NO remains unchanged.

### Chronic anemia is associated with endothelial and vascular smooth muscle dysfunction after AMI

We next investigated the effect of CA on vascular function 24 h post-AMI. Contractile responses, and endothelium-dependent and -independent relaxation responses in the isolated aortic rings using a wire myograph. Interestingly, aortic rings isolated from CA mice showed significantly reduced contractile response to phenylephrine (E_max_: 0.62 ± 0.08 mN, p=0.0002) compared to respective non-anemic mice (E_max_: 3.56 ± 0.60 mN) 24 h post-AMI (Suppl. Fig. 2, Suppl. Fig. 3A-B; Suppl. Table 4). In contrast to the AA group, the relaxation responses to acetylcholine in the presence of indomethacin were not altered in the aorta of CA mice compared to the corresponding non-anemic group (Fig. 1J; Suppl. Table 4). In the presence of L-NAME, the relaxation responses were inhibited and to a similar extent in both groups (Fig. 1K). Additionally, smooth muscle sensitivity to NO was similar in both groups (Fig. 1L; Suppl. Table 4).

We also assessed the vascular function in the femoral and saphenous arteries of the CA mice 24 h post-AMI. In contrast to the aorta, the contractile responses of femoral arteries were unchanged in CA mice compared to the respective non-anemic group (Suppl. Fig. 3C-D; Suppl. Table 5). The relaxation responses to acetylcholine in the presence of indomethacin did not differ between CA mice and non-anemic mice 24 h post-AMI (Fig. 1M; Suppl. Table 5). In the presence of L-NAME, the CA mice showed significantly reduced relaxation responses (E_max_: 73.84 ± 8.60%; p=0.04) compared to non-anemic mice (E_max_: 93.81 ± 1.48%) 24 h post-AMI (Fig. 1N; Suppl. Table 5). However, in the small resistance saphenous artery, similar to the aorta, the contractile responses to phenylephrine were significantly reduced in CA mice (E_max_: 6.71 ± 1.65 mN; p=0.01) compared to non-anemic mice (E_max_: 11.75 ± 0.88 mN) 24 h post-AMI (Suppl. Fig. 3E-F; Suppl. Table 6). Furthermore, endothelial-dependent relaxation responses to acetylcholine in the presence of indomethacin were significantly reduced in CA mice (E_max_: 67.93 ± 5.16%; p=0.004) compared to non-anemic mice (E_max_: 86.70 ± 2.95%) 24 h post-AMI (Fig. 1P; Suppl. Table 6). The SMC sensitivity to NO was preserved in the aorta of both groups (Fig. 1O & R; Suppl. Tables 5-6). However, the smooth muscle sensitivity to SNP were significantly reduced in femoral and saphenous arteries (Fig. 1O & R; Suppl. Tables 5-6). These results demonstrate that contractile responses are significantly decreased in large and small resistance arteries. Additionally, small resistance arteries also show impaired endothelial-dependent and -independent NO-mediated relaxation responses.

### Flow-mediated dilation (FMD) responses are impaired in both acute and chronic anemic mice after AMI

In addition to the detailed *ex vivo* assessment of vascular function in isolated arteries, we also assessed *in vivo* endothelial function by measuring FMD. AA mice showed significantly reduced FMD responses (5.702 ± 0.845%) compared to the respective non-anemic control group (9.270 ± 0.938%) 24 h post-AMI (Fig. 2A-B). Interestingly, similar to AA mice, CA mice also showed significantly reduced FMD responses (5.231 ± 0.547%) compared to the respective non-anemic group of mice (8.553 ± 0.767%) 24 h post-AMI (Fig. 2C-D).

**Figure 2:**
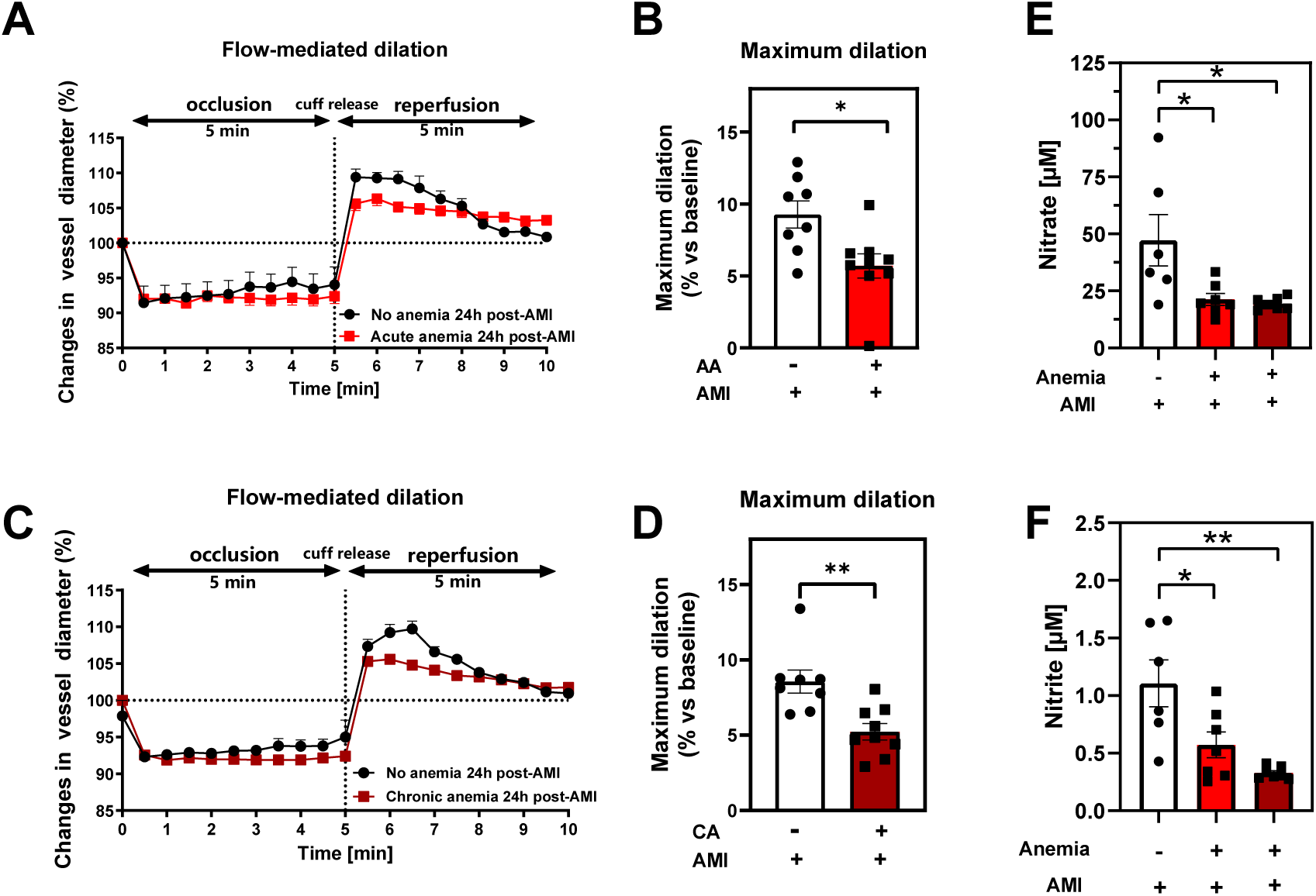
Acute and chronic anemia are associated with reduced flow-mediated dilation responses and reduced oxidative NO products 24 h post-AMI. (A, **C)** Changes in vessel diameter in acute (red squares), chronic (dark red squares), and respective non-anemic group (black circles) of mice 24 h post IR. **(B, D)** Maximal FMD response (% ratio vs. baseline). **(E, F)** Plasma nitrate and nitrite levels in the in acute (red bar), chronic (dark red bar), and respective non-anemic group (white bar). The values are presented as means ± SEM (n=8-9 per group). *, p ≤ 0.05; **, p ≤ 0.01. The average FMD in reperfusion phase was compared with the student t-test between the groups.

Endothelial dysfunction is often associated with reduced plasma oxidative nitric oxide (NOx) products. We assessed the NOx products nitrite and nitrate in the plasma. Both AA and CA mice showed significantly reduced plasma nitrate (Fig. 2E) and nitrite (Fig. 2F) levels compared to the respective non-anemic groups.

These results clearly demonstrate that, in line with the reduced *ex vivo* endothelial-dependent relaxation responses and *in vivo* FMD responses, NO bioavailability is significantly reduced in both AA and CA mice compared to respective non-anemic mice.

### Both acute and chronic anemia are associated with increased oxidative stress in the vessels

In our recent study, we showed that CA is associated with increased oxidative stress and endothelial activation (*19*). We further investigated the potential role of inflammation and reactive oxygen species (ROS) in the observed vascular dysfunction in both acute and chronic blood-loss anemia 24 h post-AMI. To detect ROS in the vessels, we performed immunofluorescence staining for the ROS product 4-HNE. The aortic sections from both AA and CA mice 24 h post-AMI showed significantly higher 4-HNE levels compared to the non-anemic mice (Fig. 3A-B). It is well known that ROS and inflammation concur with each other, so we also measured endothelial activation markers ICAM-1 and VCAM-1 in plasma. VCAM-1 was significantly increased in both AA and CA mice compared to their respective non-anemic mice 24 h post-AMI (Fig. 3C-D). These results suggest that AA and CA are associated with increased oxidative stress and endothelial inflammation after AMI.

**Figure 3:**
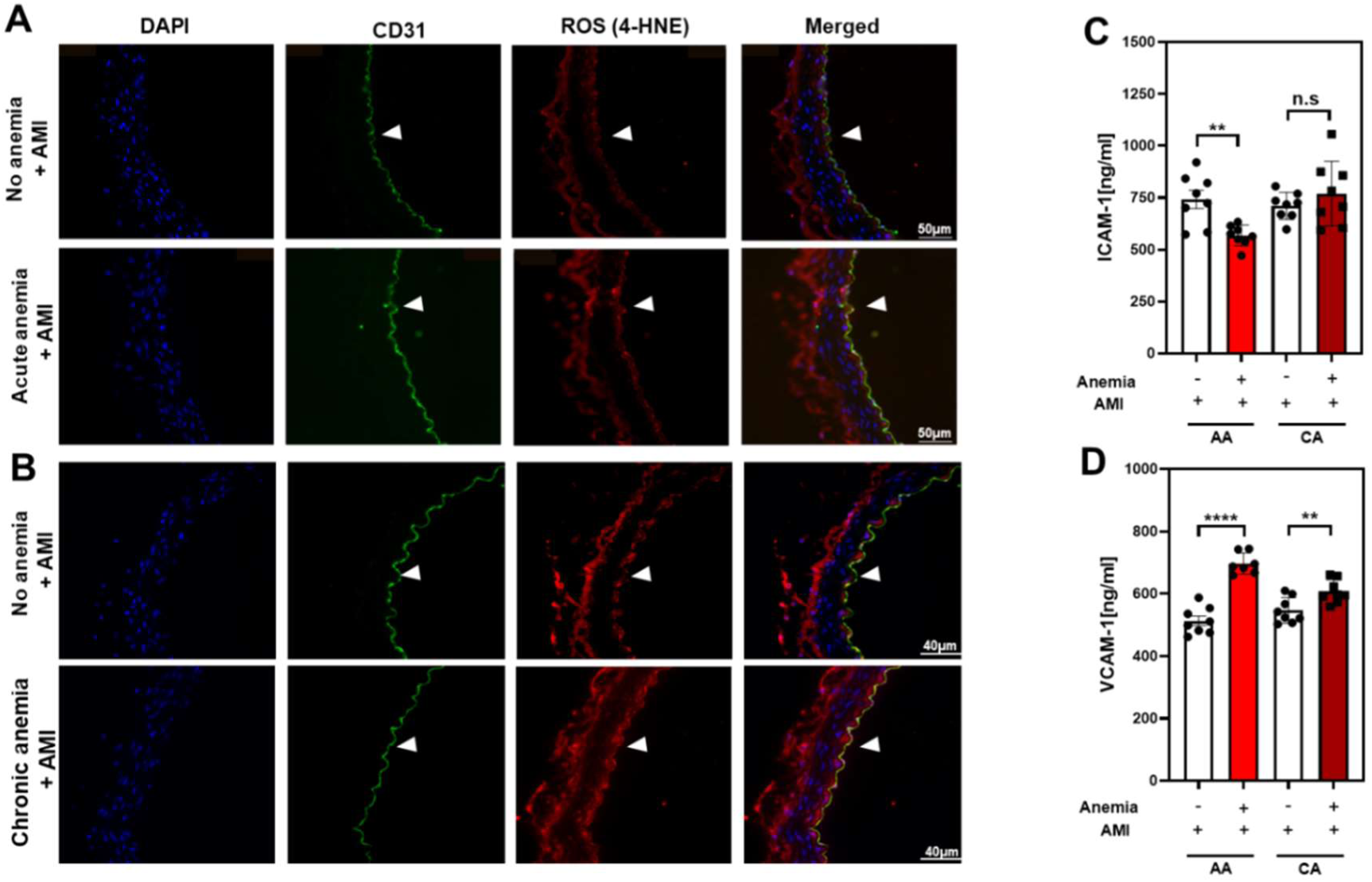
Anemia 24 h post-AMI is associated with increased oxidative stress and inflammation. Thoracic aortas isolated from acute, chronic and respective non-anemic mice 24 h post-AMI**. (A, B)** Representative images of sections that are stained with endothelial cell marker CD31 (green) and ROS-marker 4-HNE (red). DAPI (blue) staining is used to detect the nucleus. Arrowheads are in the luminal side pointing towards the respective staining. Results are representative pictures of 3 (non-anemic), 3 (acute anemic), 5 (chronic anemic) independent experiments **(C-D).** Plasma ICAM-1 and VCAM-1 levels in acute (AA, red bar), chronic (CA, dark red bar), and respective non-anemic group (white bar) of mice 24 h post-AMI. All values are presented as means ± SEM. **, p ≤ 0.01; ****, p ≤ 0.0001; ns, not significant.

### NAC treatment reversed the vascular dysfunction in both acute and chronic anemic mice

From our data, it is evident that ROS mediates vascular dysfunction in both AA and CA mice. To further confirm this, we supplemented mice with NAC and assessed vascular function after AMI. The contractile responses to phenylephrine were not altered between AA mice and the respective non-anemic group of mice (Suppl. Table 1). Interestingly, after NAC supplementation, the abrogated contractile responses in CA mice were significantly improved in the aorta (E_max_: 3.14 ± 0.64 mN; p=0.004) compared to CA mice without NAC treatment (E_max_: 0.62 ± 0.08 mN) 24 h post-AMI (Suppl. Fig. 3G-H; Suppl. Table 4). The contractile responses were similar between CA mice and non-anemic mice 24 h post-AMI. Similar improvements in contractile responses were also observed in saphenous arteries (Suppl. Fig. 3K-L). Next, in the same mice, we assessed endothelium-dependent and -independent relaxation responses in both AA and CA mice and the respective control group 24 h post-AMI. As expected, in both anemic groups and all types of arteries, the endothelium-dependent relaxation responses were improved with NAC treatment (Fig. 4A, D, G, J, M&P Suppl. Fig. 4A-F; Suppl. Tables 1-6). The endothelium-independent relaxation to SNP was preserved in both anemic groups and all three types of arteries (Fig. 4C, F, I, L, O&R; Suppl. Tables 1-6).

**Figure 4:**
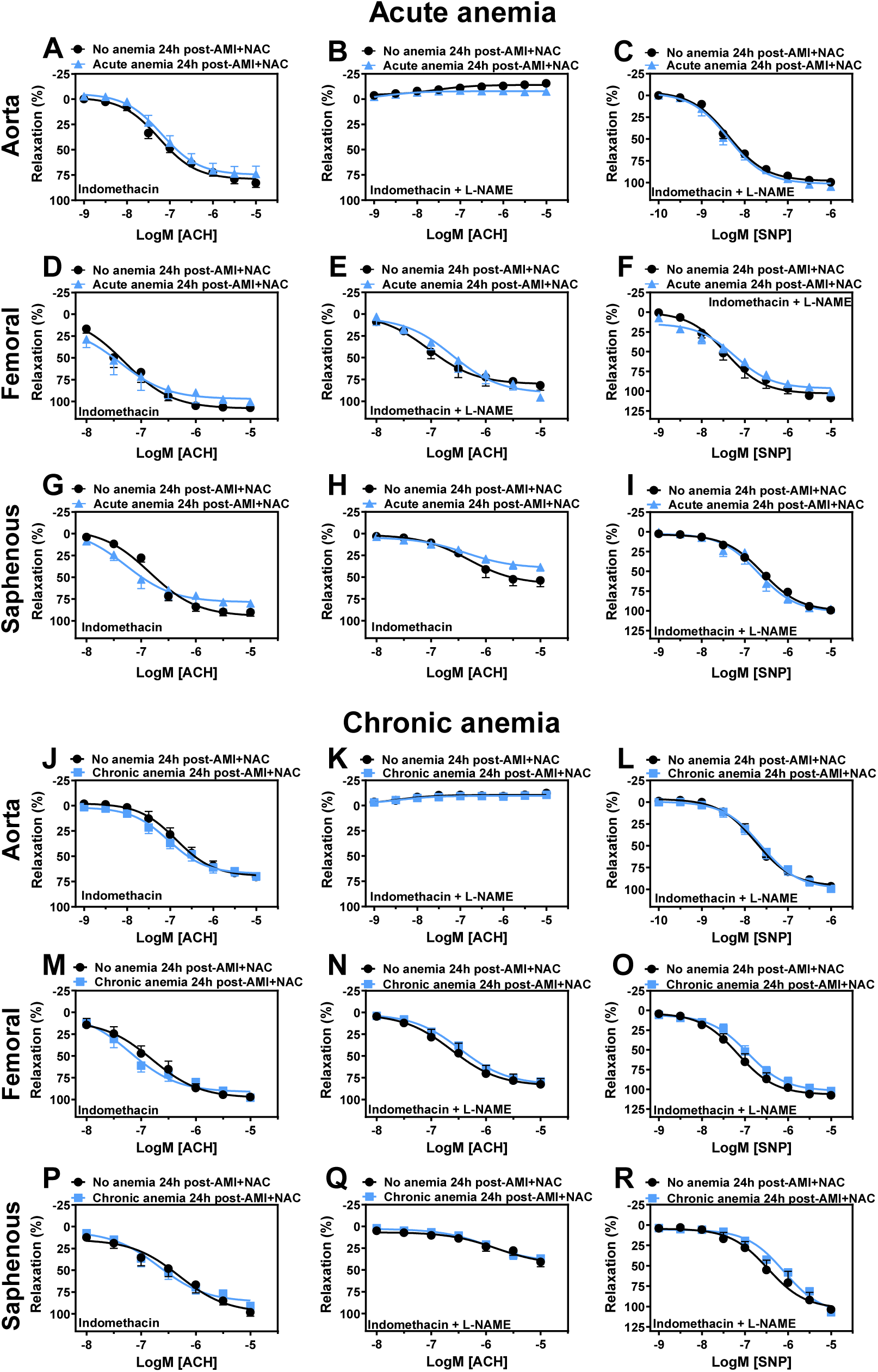
NAC supplementation improves endothelial-dependent relaxation response in anemia 24 h post-AMI. Arterial segments of aorta **(A-C, J-L)**, femoral **(D-F, M-O)** and saphenous arteries **(G-I, P-R)** were isolated from acute (light blue triangles) and chronic (light blue squares) anemic mice and respective non-anemic mice (black circles) 24h post-AMI. Arterial segments were pre-contracted using phenylephrine (PHE; 10 µM), and their relaxation responses to acetylcholine (ACH; 1 nM-10 µM) were assessed using wire myography. **(A, D, G, J, M, P)** Relaxation (%) in the presence of indomethacin (10 µM, a COX inhibitor). **(B, E, H, K, N, Q)** Relaxation in the presence of indomethacin and L-NAME (100 µM, NOS inhibitor). Maximal endothelial-dependent relaxation response (E_max, %_). **(C, F, I, L, O, R)** Relaxation responses to sodium nitroprusside (SNP, 10 nM-10 µM) in the presence of indomethacin and L-NAME. All values are mean values ± SEM (n = 8–10 per group). Concentration-response curves (CRCs) were analysed by Two-Way ANOVA and Bonferroni ‘s post-hoc test to compare acute, chronic and respective sham group of mice.

These results demonstrate that NAC treatment effectively reverses vascular dysfunction in both acute and CA mice by improving both contractile and endothelium-dependent relaxation responses, indicating the critical role of ROS in mediating vascular dysfunction.

### NAC supplementation improved flow-mediated dilation (FMD) responses in both acute and chronic anemic mice after AMI

We further assessed the FMD responses in both AA and CA mice and their respective non-anemic groups 24 h post-AMI. In line with *ex vivo* data, after NAC supplementation, both AA and CA mice showed improved FMD responses (Fig. 5A-D; Suppl. Fig. 5A-D). This further confirms that ROS mediates vascular dysfunction in both anemic groups after AMI. Additionally, after NAC supplementation, both nitrite and nitrate levels improved in AA and CA mice compared to the respective non-anemic groups 24 h post-AMI (Fig. 5E-F). These results conclude that NAC supplementation improves endothelial dysfunction and the bioavailability of NO.

**Figure 5:**
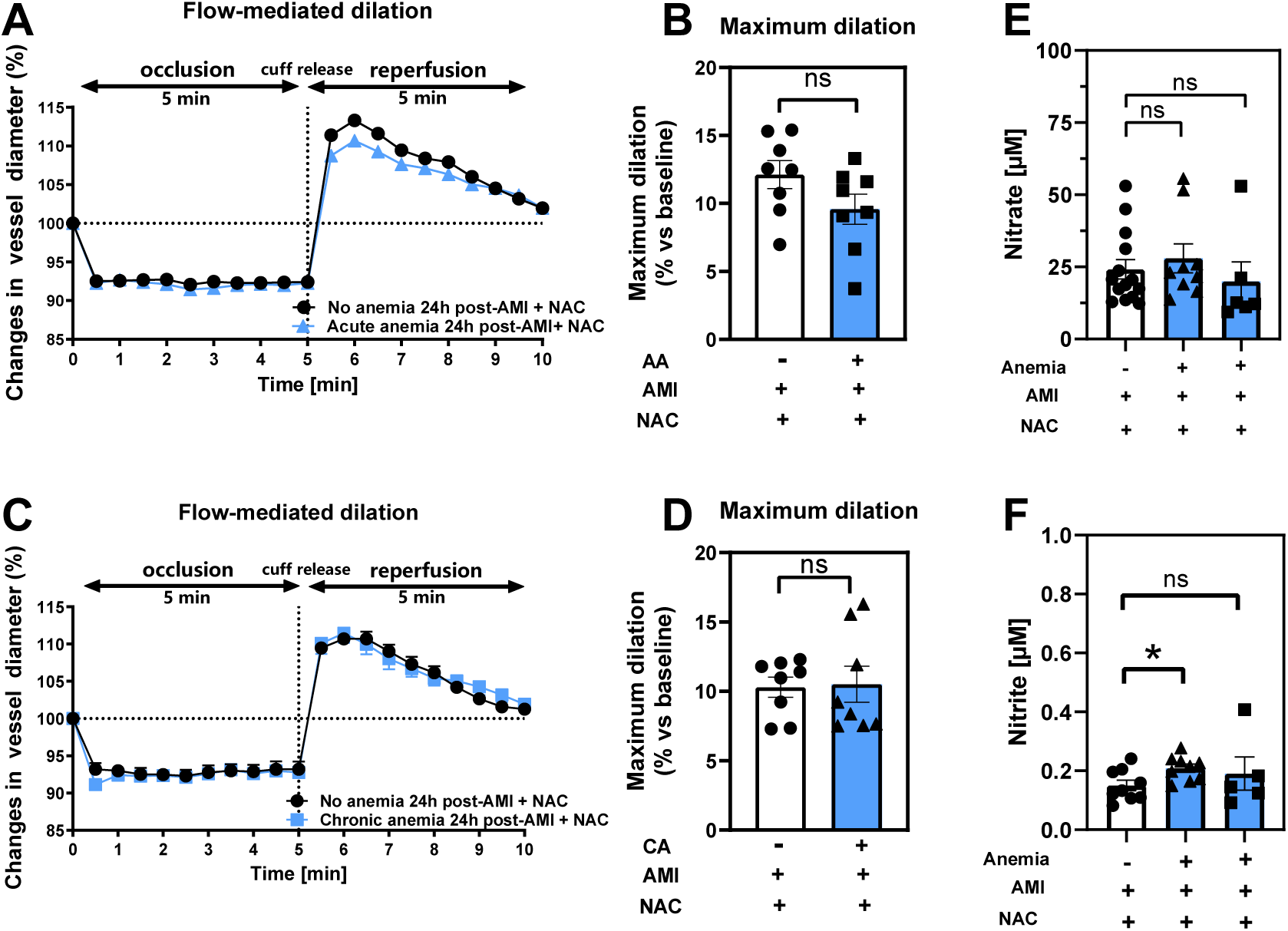
NAC treatment improves flow-mediated dilation (FMD) response in acute and chronic anemia 24 h post-AMI. (A, C) Changes in vessel diameter in acute (light blue triangles), chronic (light blue, squares), and respective non-anemic groups (black, circles) of mice 24 h post-AMI. (B, D) Maximal FMD response (% ratio vs. baseline). **(E, F)** Plasma nitrate and nitrite levels in the in acute (light blue bar, triangles), chronic (light blue bar, squares), and respective non-anemic group (white bar, circles). The values are presented as means ± SEM (n=8-9 per group). *, p ≤ 0.05; **, p ≤ 0.01; ns, not significant. The average FMD in the reperfusion phase was compared with the Student’s t-test between the groups.

By performing immunofluorescence staining, we demonstrated that NAC-treated mice had reduced ROS formation in the aortic segments (Fig. 6A-B). Additionally, the increased plasma levels of VCAM-1 were normalized in the CA mice after NAC treatment (Fig. 3D and 6C) whereas they remained high in AA mice (Fig. 3D and 6C). These findings support a potential role for ROS and inflammation in anemia-associated endothelial dysfunction post-AMI.

**Figure 6:**
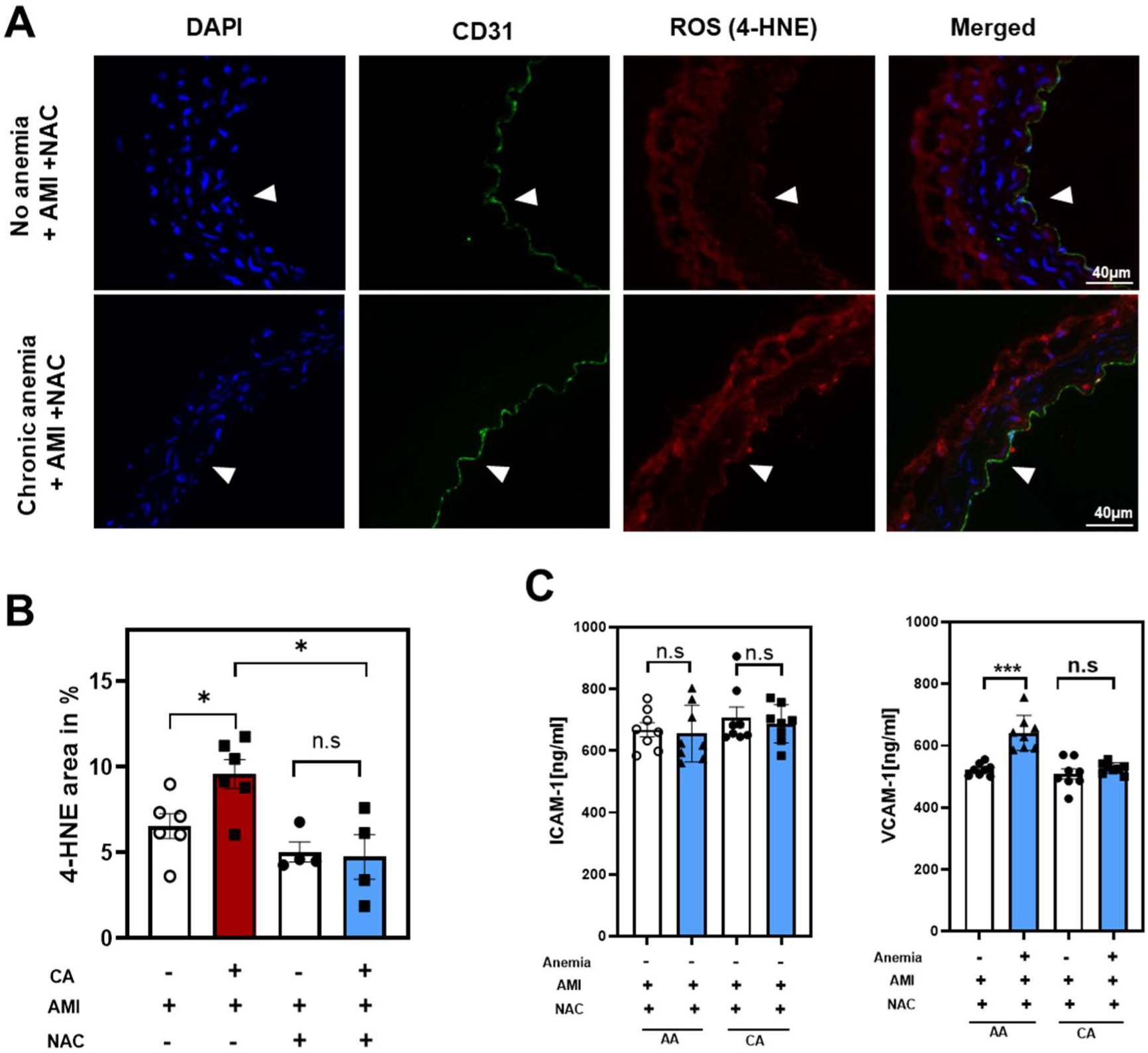
NAC treatment reduces ROS formation and endothelial inflammation in anemic mice 24 h post-AMI. **(A-B)** Thoracic aortas isolated from chronic anemic mice and respective non-anemic mice 24 h post IR. **(A)** Representative images of sections that are stained with endothelial cell marker CD31 (green) and ROS-marker 4-HNE (red). DAPI (blue) staining is used to detect the nucleus. Arrowheads are in the luminal side pointing towards the respective staining. For statistical analysis of the co-staining six images were evaluated and were compared using a student t-test between sham and anemic groups. Scale bar=40 µm. **(B)** Quantification of 4-HNE in chronic (dark red bar, squares), non-anemic (white bar, open circles), NAC treated chronic anemic (blue bar, squares) and non-anemic (white bar, closed circles) group of mice 24 h post-AMI. **(C)** Plasma ICAM-1 and VCAM-1 levels in acute (light blue, triangles), chronic (light blue, squares), and respective non-anemic groups (white bar, open and closed circles) of mice 24 h post-AMI. All values are presented as means ± SEM. *, p ≤ 0.05; ***, p ≤ 0.001; ns, not significant.

### Anemic RBCs promote endothelial dysfunction

Previous studies have demonstrated that RBCs induce endothelial dysfunction in various cardiometabolic diseases (*20*). Given this, we investigated the potential role of anemic RBCs in mediating ED after AMI. First, we incubated RBCs from anemic and non-anemic mice 24 h post-AMI, co-incubated (2 h at 37°C) with WT aortic rings, and assessed the vascular responses. The endothelium-dependent relaxation responses to acetylcholine in the presence of indomethacin were completely inhibited in the aortic rings incubated with anemic RBCs compared to non-anemic mice (Figure 7A). The relaxation responses in the presence of L-NAME were completely inhibited in both groups (Figure 7B), indicating that relaxation responses are entirely mediated by NO in these vessels. In addition, the relaxation responses to SNP were mildly abrogated in anemic RBCs-incubated aortic rings compared to the non-anemic group (Figure 7C).

**Figure 7:**
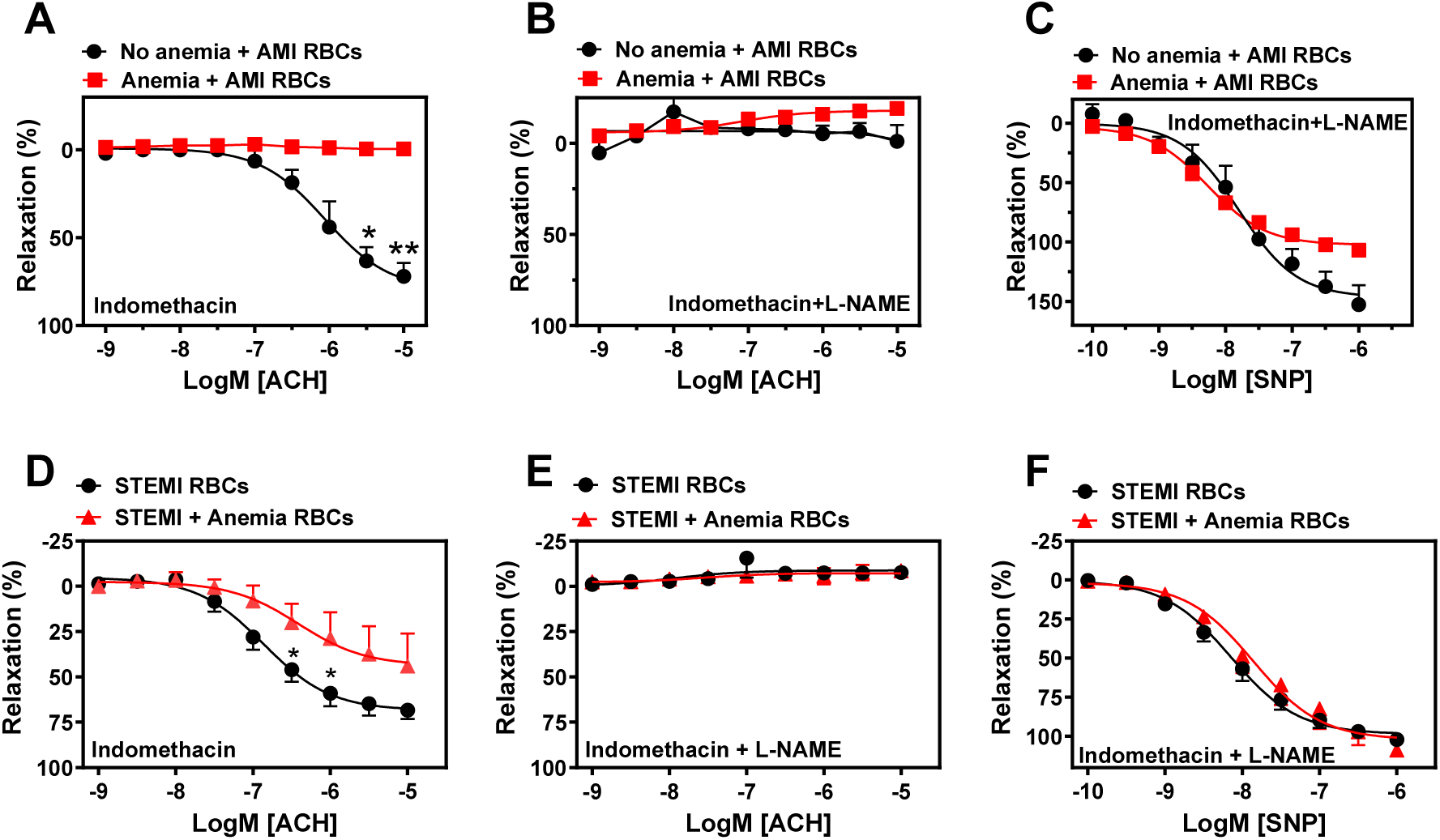
RBCs from anemic mice 24h post-AMI and anemic STEMI patients induce endothelial dysfunction. **(A-D)** RBCs were isolated from non-anemic (black circles) and anemic mice (red squares) 24 h post-AMI. **(E-H)** RBCs were isolated from STEMI patients without anemia (black circles) and with anemia (red triangles). Haematocrit (40%) was prepared in a KREBS buffer. Aortic rings from WT mice were incubated with haematocrit either 2 (mice) or 6 (humans) h and mounted in wire myograph. **(B, F)** Relaxation responses (%) to acetylcholine in the presence of indomethacin (10 µM, COX inhibitor) in aortic rings incubated with mice RBCs. **(C, G**) Relaxation responses to acetylcholine in the presence of indomethacin (10 µM, COX inhibitor). **(D, H)** Relaxation responses to sodium nitroprusside (SNP, 10 nM-10 µM) in the presence of indomethacin and L-NAME. Values are shown as means ± SEM. n= 4 for mice (per group). STEMI group: n=8, STEMI + anemia: n=4. *, p ≤ 0.05; **, p ≤ 0.01. CRCs were analysed by Two-Way ANOVA and Bonferroni ‘s post-hoc test.

We also investigated whether RBCs from anemic STEMI patients induce ED. Our results showed that aortic rings incubated with RBCs from anemic STEMI patients showed reduced endothelial-dependent relaxation responses compared to STEMI patients without anemia (Figure 7D). These relaxation responses are endothelial-dependent and mediated by NO (Figure 7E). Endothelium-independent relaxation responses to SNP were similar in both groups (Figure 7F). These results further demonstrate that RBCs from anemic STEMI patients induce ED by altering the NO-mediated relaxation responses.

## Discussion

In this study, we examined the effect of acute and chronic anemia on vascular function after AMI. The key findings of the study are that (1) AA and CA are associated with reduced endothelial NO-dependent relaxation responses *ex vivo* and impaired *in vivo* flow-mediated dilation responses 24 h post-AMI, (2) Plasma nitrate and nitrite levels were significantly reduced in AA and CA mice 24 h post-AMI, (3) VSMC contractile responses were specifically abrogated in large and small resistance arteries of CA mice 24 h post-AMI, (4) AA and CA are associated with increased production of reactive oxygen species (ROS) in the vessels, (5) NAC treatment improved vascular function (both contractile and relaxation responses) in AA and CA mice 24 h post-AMI, (6) RBCs from anemic mice 24 h post-AMI and STEMI patients with anemia induced vascular dysfunction in murine aortic rings, highlighting the potential role of RBCs in endothelial dysfunction.

Several comorbidities lead to adverse outcomes after AMI, particularly anemia, which is clinically presented in elderly patients during admission, hospitalization, or post-AMI (*27, 4*). Understanding the effects of blood loss anemia on vascular function is clinically relevant, as it is the main cause of reduced Hb during hospitalization due to blood sampling for laboratory tests or clinical interventions such as the insertion of catheters, endotracheal tubes, etc. (*28*). These interventions cannot be avoided in the hospital setting. Therefore, in this study, we examined both acute and chronic blood loss anemic mouse models, which reflect blood loss anemia during hospitalization in patients. Of note, we previously showed that these mouse models are mildly anemic, and we did not observe any severe consequences due to blood loss, such as oedema (*18, 19*). Additionally, AMI was induced in both mouse models to investigate the possible reasons for the poor prognosis of anemic patients after AMI in relation to overall vascular function. Interestingly, our findings demonstrate that AA results in a worsening of endothelial function in different types of vessels and also *in vivo* FMD responses.

In our previous study we demonstrated that AA (without AMI) is associated with compensatory increased FMD responses (*18*). However, after AMI, the compensatory improved FMD responses are compromised, which is explained by increased oxidative stress and endothelial activation/inflammation. In addition, previous clinical studies demonstrated that anemia without comorbidities results in increased FMD responses in healthy volunteers, whereas anemia in combination with CKD or diabetes worsens endothelial function and thus FMD responses (*14–16*). Altogether, co-morbidities and acute stress condition such as AMI worsen the endothelial function in anemia. To our knowledge, this is the first study to demonstrate how endothelial function is affected after AMI in anemia.

Studies have shown that uncoupling of eNOS leads to ED in several pathologies (*29*). Additionally, in different disease states, it has been shown under oxidative stress conditions, O_2_ reacts with NO to form ONOO^-^, resulting in eNOS uncoupling, lipid peroxidation, and vascular damage (*30, 31*). Moreover, eNOS uncoupling leads to generation superoxide instead of NO, further becoming a source of detrimental free radicals, impairing endothelial function (*32*). In this study, we observed an increase in the ROS oxidative product 4-HNE in the vessels of both AA and CA mice, indicating the potential role of ROS in mediating detrimental effects on the endothelium and, consequently, on NO bioavailability. In our recent study, we demonstrated that endothelium-dependent relaxation responses are impaired in aortic rings from CA mice (*19*). However, 24 h post-AMI, we did not observe significantly impaired relaxation responses in CA mice compared to non-anemic mice. A possible explanation for this could be that the amplitude of contraction of VSMC was drastically reduced, which may have masked the changes in endothelium-dependent relaxation measurements to acetylcholine in the *ex vivo* setting. The relaxation and contractile responses in femoral arteries are preserved in CA mice compared to the respective sham group. However, in small resistance saphenous arteries, the contraction and relaxation responses are significantly impaired in CA mice after AMI. These results indicate that the effects of CA vary across different vascular beds following AMI.

The effect of anemia on VSMC dysfunction is not well studied. However, a recent study demonstrated that intracellular iron deficiency leads to VSMC dysfunction, resulting in pulmonary hypertension mediated by increased endothelin (*33*). In addition, sickle cell anemia is also associated with VSMC dysfunction in pulmonary circulation due to increased oxidative stress, severe haemolysis, and inflammation (*34*). In the current study, we demonstrated that AA does not affect SMC contractile responses, whereas these responses are abrogated in CA. We believe that these effects are caused by chronic mild hypoxia due to the diminished oxygen-carrying capacity of RBCs to vascular smooth muscle cells in CA mice (*35*). The other possible explanation is that chronic hypoxia results in increased ROS production, which might disrupt the contractile machinery in VSMCs (*36*). However, we do not know the exact mechanism of how anemic RBCs impair the contractile responses which will be evaluated in the future.

Recent studies from our group and others demonstrated that RBCs plays a crucial role in cardiovascular diseases. In addition, Several studies demonstrated that RBCs mediate ED in various pathological conditions, such as diabetes mellitus and hypercholesterolemia, due to elevated arginase 1 levels and increased ROS formation in endothelial cells mediated by RBC (*21, 22*). Our results from the co-incubation experiments of RBCs from anemic mice 24 h post-AMI and anemic STEMI patients with murine aortic rings show a clear induction of ED in the aortic rings. However, performing co-incubation studies with RBCs and Nω-Hydroxy-nor-L-arginine acetate (nor-NOHA), a specific arginase inhibitor, did not show any improvement in endothelial function after arginase inhibition in the murine aortas (data not shown). These results suggest a different underlying mechanism of inducing ED in anemia. Extracellular vesicles (EVs) are known to mediate intercellular communication by transferring different molecules and therefore playing an important role in cardio vascular diseases (*37, 38*). Previous studies in sickle cell anemia and diabetes mellitus type 2 have implicated RBC-derived extracellular vesicles (REVs) in mediating vascular dysfunction (*39, 40*). In the current study, we primarily focused on the assessment and comparison of vascular function in acute and chronic anemia after AMI. The investigation of the potential role of REVs in anemia-associated ED after AMI was beyond the scope of this study, but it needs further evaluation. Notably, we cannot overlook some limitations of this study. We strongly believe that the observations made here need to be re-evaluated in another type of anemic mouse model to determine whether the effects are specific to this particular type of anemia or apply to all types of anemia.

### Conclusions

Our data suggest that both acute and chronic blood-loss anemia are associated with decreased NO production after AMI due to increased inflammation and ROS production. Additionally, chronic anemia specifically led to reduced smooth muscle contractile responses. We also demonstrated that RBCs from anemic STEMI patients attenuate endothelial-dependent relaxation responses as well as vascular smooth muscle contractile responses, highlighting the potential role of anemic RBCs in vascular dysfunction. Furthermore, we showed that treating anemic mice with a ROS scavenger (NAC) improved relaxation responses in both acute and chronic anemic mice. Taken together, NAC supplementation improves vascular function in blood-loss anemia after AMI.

## Data availability statement

The raw data supporting the conclusions of this manuscript will be made available by the authors, without undue reservation.

## Author Contributions

IS and RC designed the work. IS contributed to qRT-PCR, biochemical assays, Western Blot analysis. IS, AS, FC, RC, VY contributed to myograph and organ bath experiments. PW contributed to arrangement of STEMI patient samples. AS, IS contributed to immunohistochemistry. SB contributed to anemia induction. IS and RC interpreted the data and drafted the manuscript. MK, CJ, NG, RC obtained the funding. MRH, AP, CJ, NG, MK, RC critically revised the manuscript. All authors contributed to the article and approved the data for submission.

## Acknowledgments

The authors wish to thank Dr. Susanne Pfeiler for organisational help.

