## Supplementary Figures 1-7 for "Distinct effects of acute and chronic blood loss anemia on vascular function after acute myocardial infarction"

### Slide 1
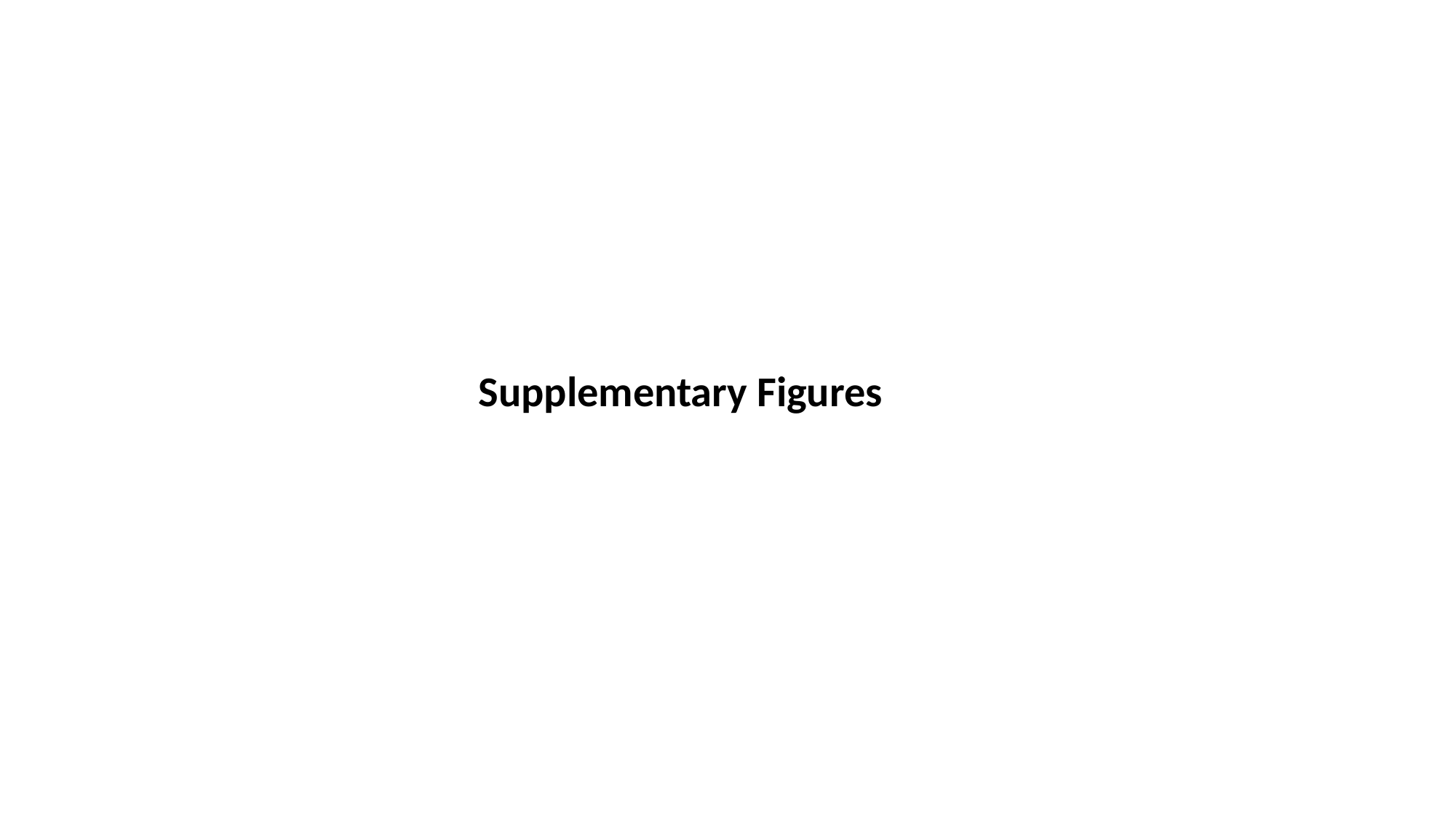

#### Supplementary Figures

### Slide 2
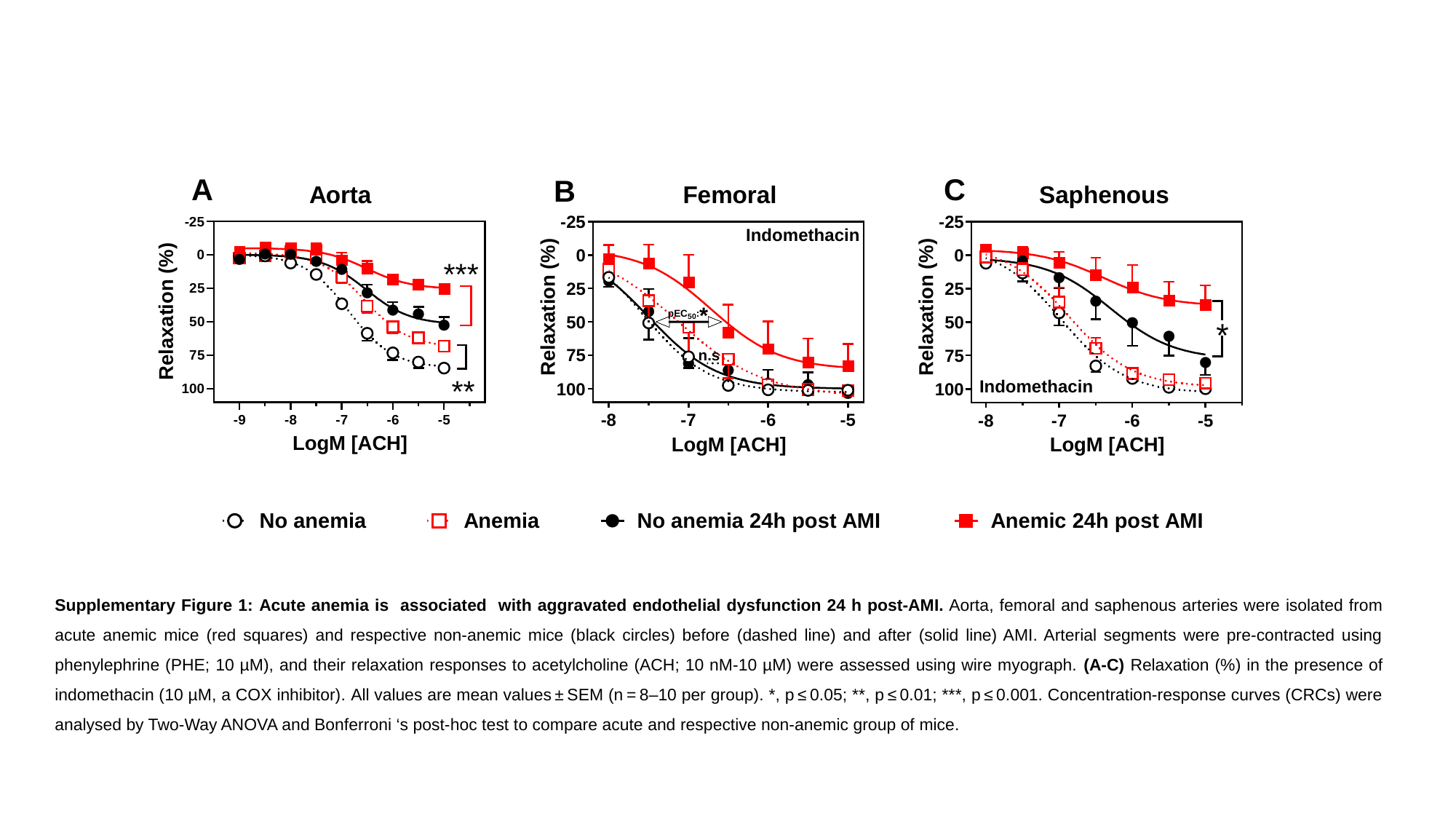

Supplementary Figure 1: Acute anemia is associated with aggravated endothelial dysfunction 24 h post-AMI. Aorta, femoral and saphenous arteries were isolated from acute anemic mice (red squares) and respective non-anemic mice (black circles) before (dashed line) and after (solid line) AMI. Arterial segments were pre-contracted using phenylephrine (PHE; 10 µM), and their relaxation responses to acetylcholine (ACH; 10 nM-10 µM) were assessed using wire myograph. (A-C) Relaxation (%) in the presence of indomethacin (10 µM, a COX inhibitor). All values are mean values ± SEM (n = 8–10 per group). *, p ≤ 0.05; **, p ≤ 0.01; ***, p ≤ 0.001. Concentration-response curves (CRCs) were analysed by Two-Way ANOVA and Bonferroni ‘s post-hoc test to compare acute and respective non-anemic group of mice.

### Slide 3
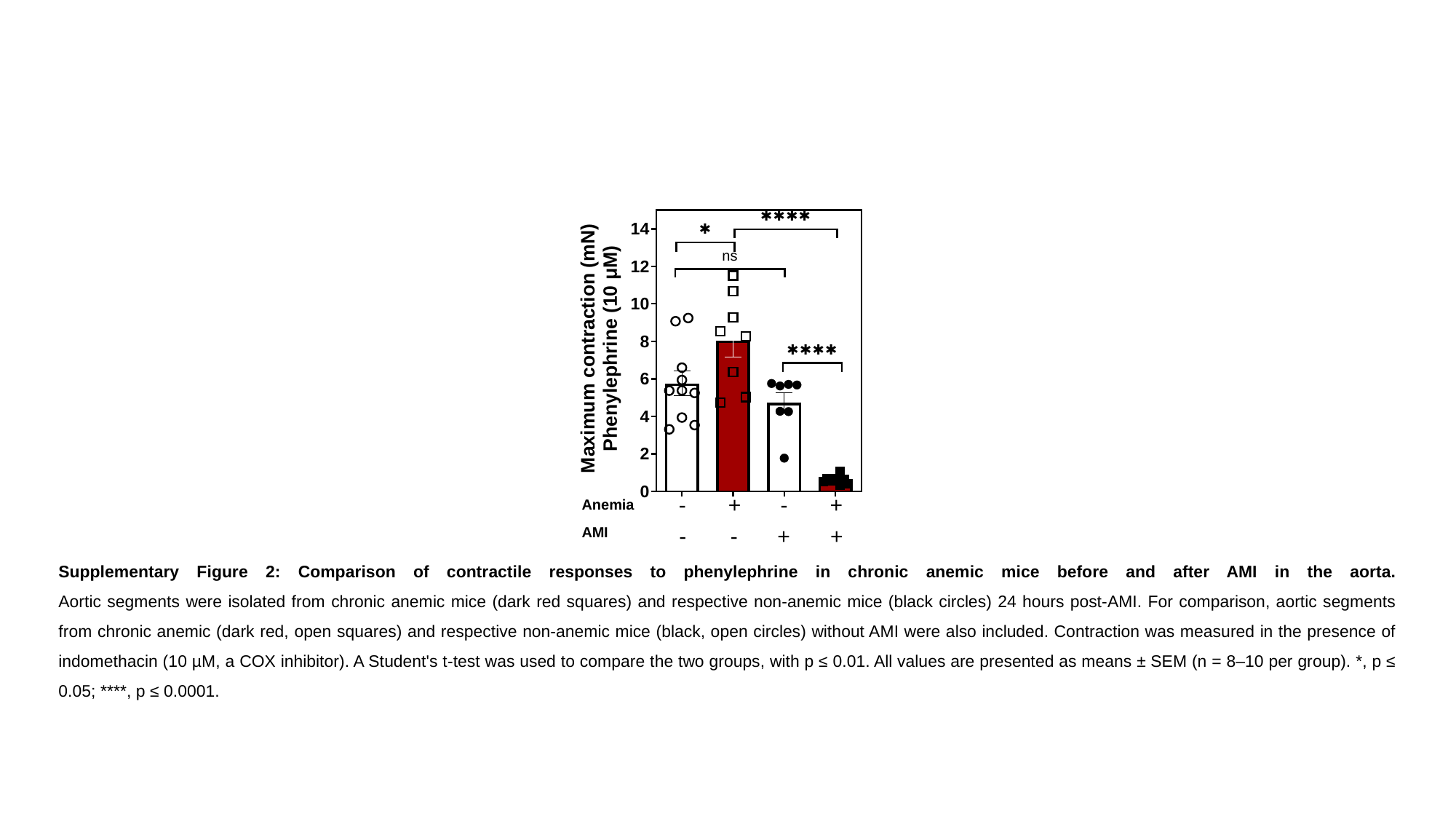

Supplementary Figure 2: Comparison of contractile responses to phenylephrine in chronic anemic mice before and after AMI in the aorta.
Aortic segments were isolated from chronic anemic mice (dark red squares) and respective non-anemic mice (black circles) 24 hours post-AMI. For comparison, aortic segments from chronic anemic (dark red, open squares) and respective non-anemic mice (black, open circles) without AMI were also included. Contraction was measured in the presence of indomethacin (10 µM, a COX inhibitor). A Student's t-test was used to compare the two groups, with p ≤ 0.01. All values are presented as means ± SEM (n = 8–10 per group). *, p ≤ 0.05; ****, p ≤ 0.0001.

### Slide 4
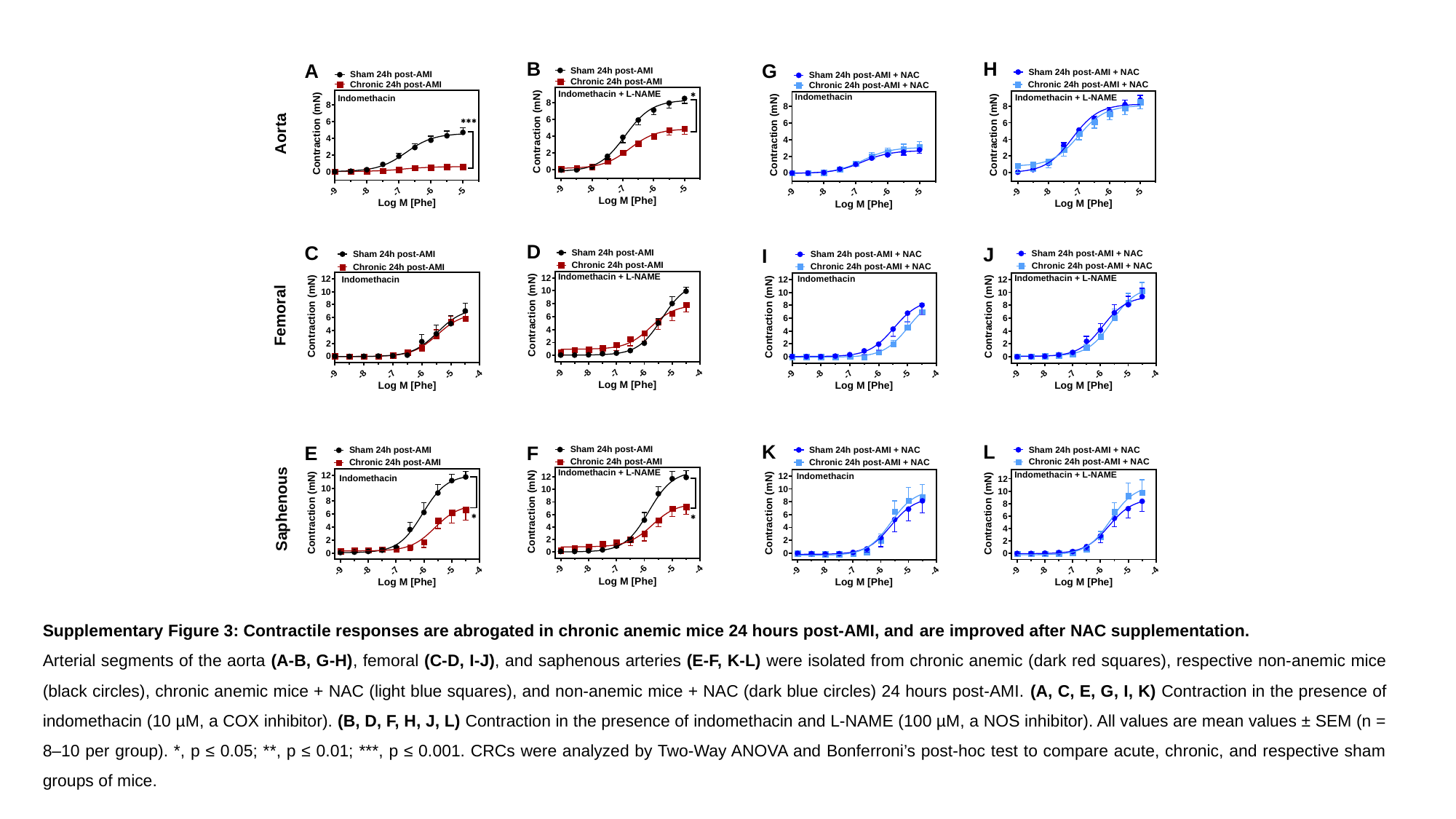

Supplementary Figure 3: Contractile responses are abrogated in chronic anemic mice 24 hours post-AMI, and are improved after NAC supplementation.
Arterial segments of the aorta (A-B, G-H), femoral (C-D, I-J), and saphenous arteries (E-F, K-L) were isolated from chronic anemic (dark red squares), respective non-anemic mice (black circles), chronic anemic mice + NAC (light blue squares), and non-anemic mice + NAC (dark blue circles) 24 hours post-AMI. (A, C, E, G, I, K) Contraction in the presence of indomethacin (10 µM, a COX inhibitor). (B, D, F, H, J, L) Contraction in the presence of indomethacin and L-NAME (100 µM, a NOS inhibitor). All values are mean values ± SEM (n = 8–10 per group). *, p ≤ 0.05; **, p ≤ 0.01; ***, p ≤ 0.001. CRCs were analyzed by Two-Way ANOVA and Bonferroni’s post-hoc test to compare acute, chronic, and respective sham groups of mice.

### Slide 5
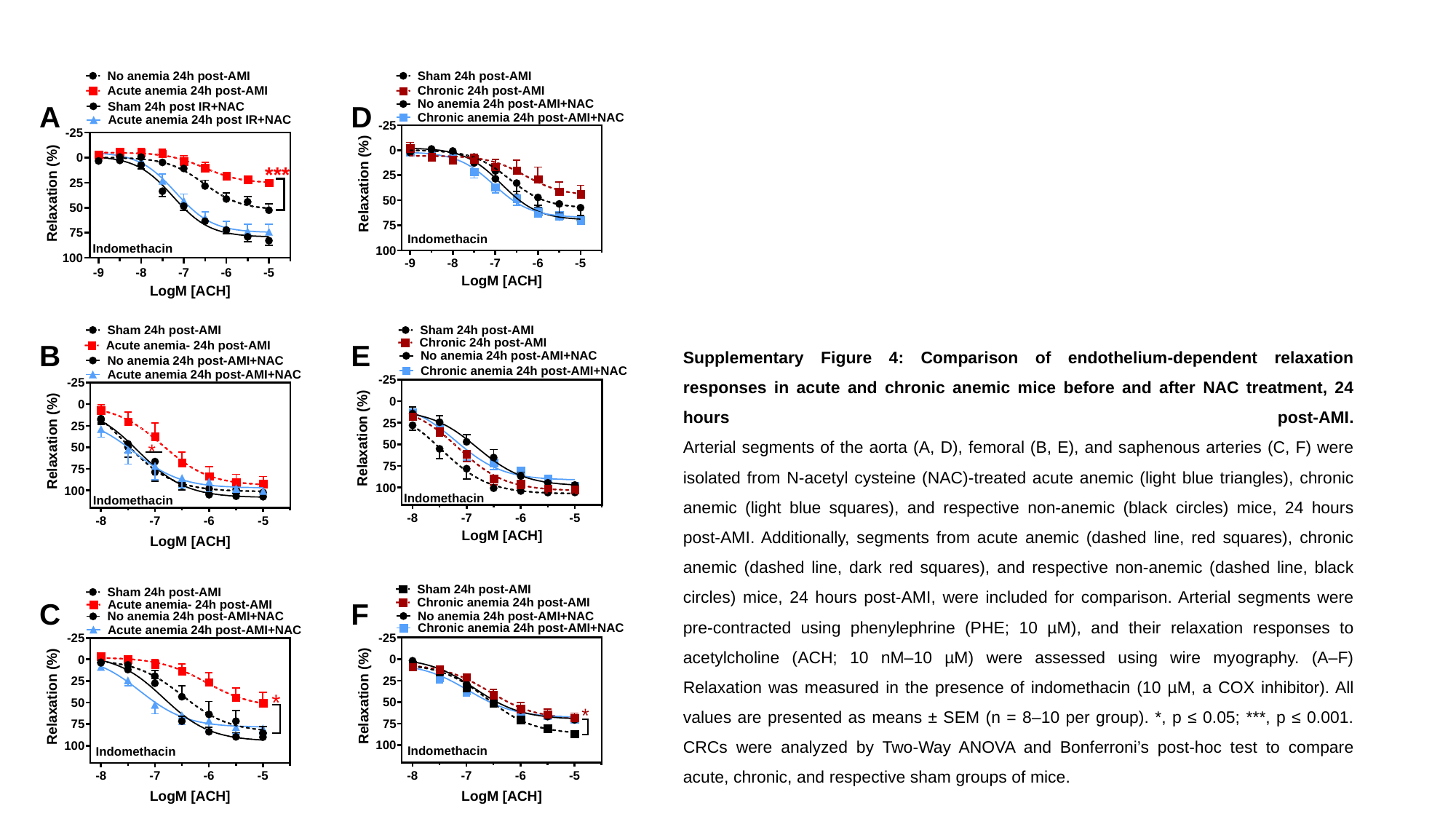

Supplementary Figure 4: Comparison of endothelium-dependent relaxation responses in acute and chronic anemic mice before and after NAC treatment, 24 hours post-AMI.
Arterial segments of the aorta (A, D), femoral (B, E), and saphenous arteries (C, F) were isolated from N-acetyl cysteine (NAC)-treated acute anemic (light blue triangles), chronic anemic (light blue squares), and respective non-anemic (black circles) mice, 24 hours post-AMI. Additionally, segments from acute anemic (dashed line, red squares), chronic anemic (dashed line, dark red squares), and respective non-anemic (dashed line, black circles) mice, 24 hours post-AMI, were included for comparison. Arterial segments were pre-contracted using phenylephrine (PHE; 10 µM), and their relaxation responses to acetylcholine (ACH; 10 nM–10 µM) were assessed using wire myography. (A–F) Relaxation was measured in the presence of indomethacin (10 µM, a COX inhibitor). All values are presented as means ± SEM (n = 8–10 per group). *, p ≤ 0.05; ***, p ≤ 0.001. CRCs were analyzed by Two-Way ANOVA and Bonferroni’s post-hoc test to compare acute, chronic, and respective sham groups of mice.

### Slide 6
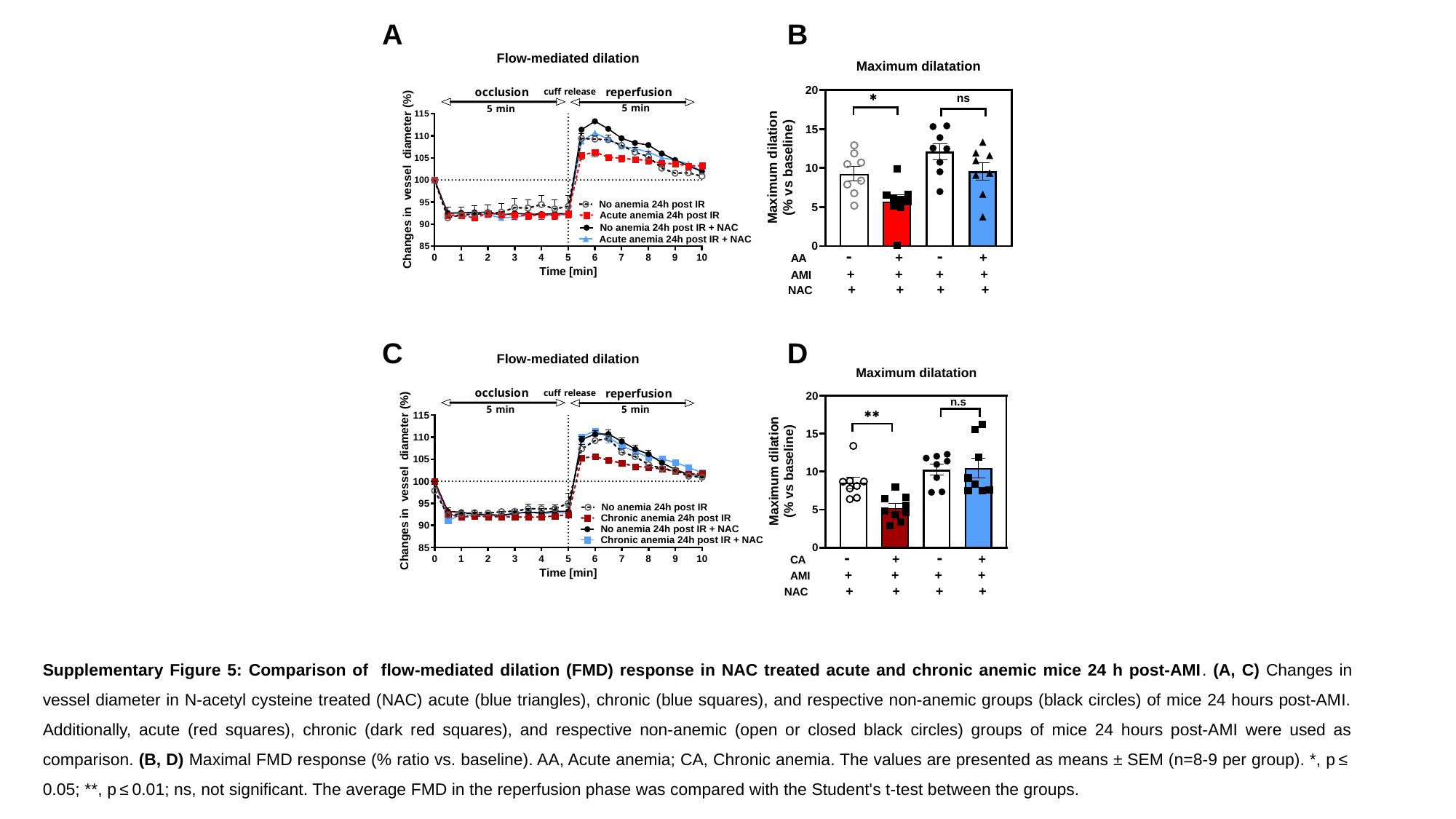

Supplementary Figure 5: Comparison of flow-mediated dilation (FMD) response in NAC treated acute and chronic anemic mice 24 h post-AMI. (A, C) Changes in vessel diameter in N-acetyl cysteine treated (NAC) acute (blue triangles), chronic (blue squares), and respective non-anemic groups (black circles) of mice 24 hours post-AMI. Additionally, acute (red squares), chronic (dark red squares), and respective non-anemic (open or closed black circles) groups of mice 24 hours post-AMI were used as comparison. (B, D) Maximal FMD response (% ratio vs. baseline). AA, Acute anemia; CA, Chronic anemia. The values are presented as means ± SEM (n=8-9 per group). *, p ≤ 0.05; **, p ≤ 0.01; ns, not significant. The average FMD in the reperfusion phase was compared with the Student's t-test between the groups.

### Slide 7
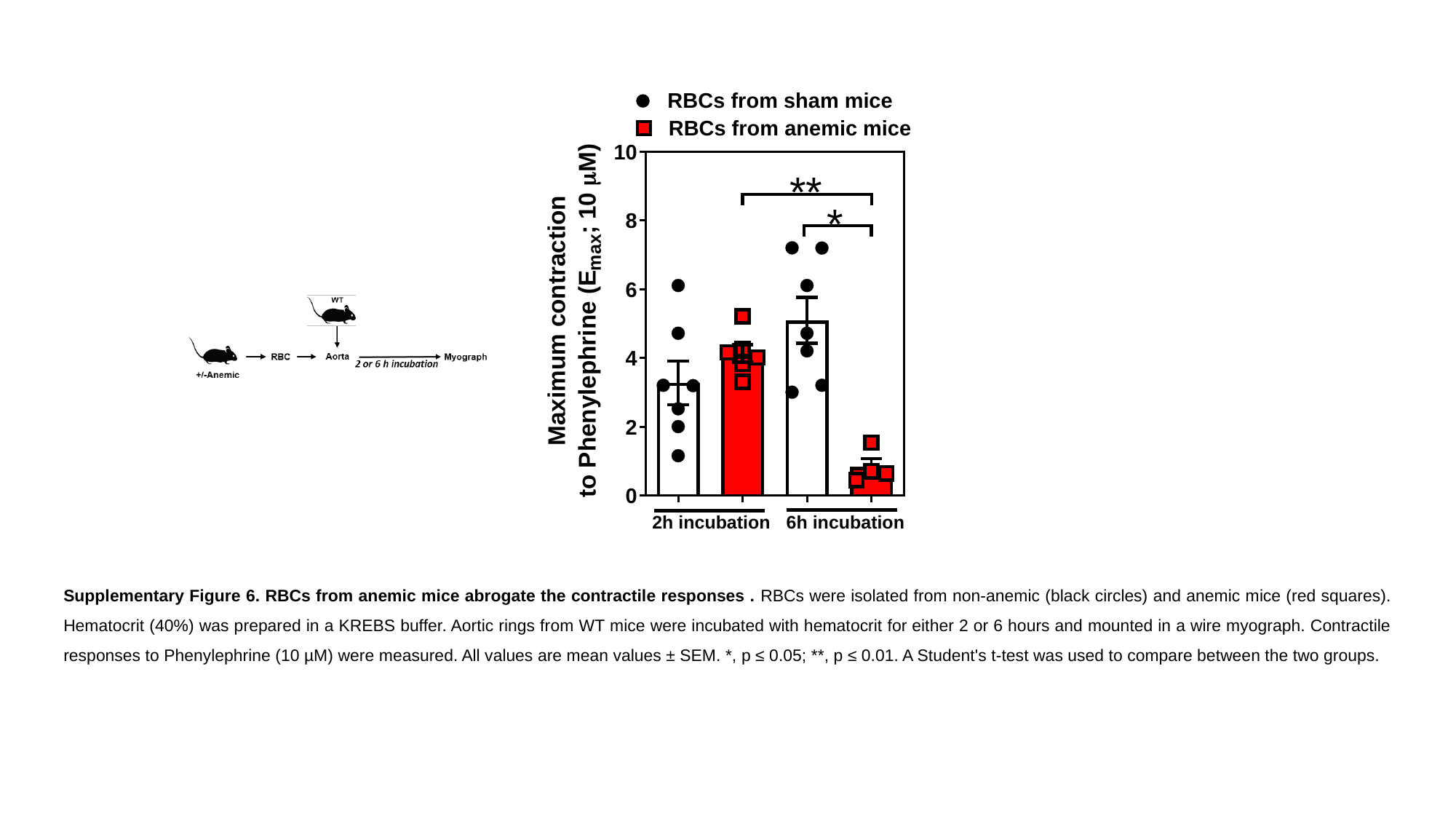

Supplementary Figure 6. RBCs from anemic mice abrogate the contractile responses . RBCs were isolated from non-anemic (black circles) and anemic mice (red squares). Hematocrit (40%) was prepared in a KREBS buffer. Aortic rings from WT mice were incubated with hematocrit for either 2 or 6 hours and mounted in a wire myograph. Contractile responses to Phenylephrine (10 µM) were measured. All values are mean values ± SEM. *, p ≤ 0.05; **, p ≤ 0.01. A Student's t-test was used to compare between the two groups.

### Slide 8
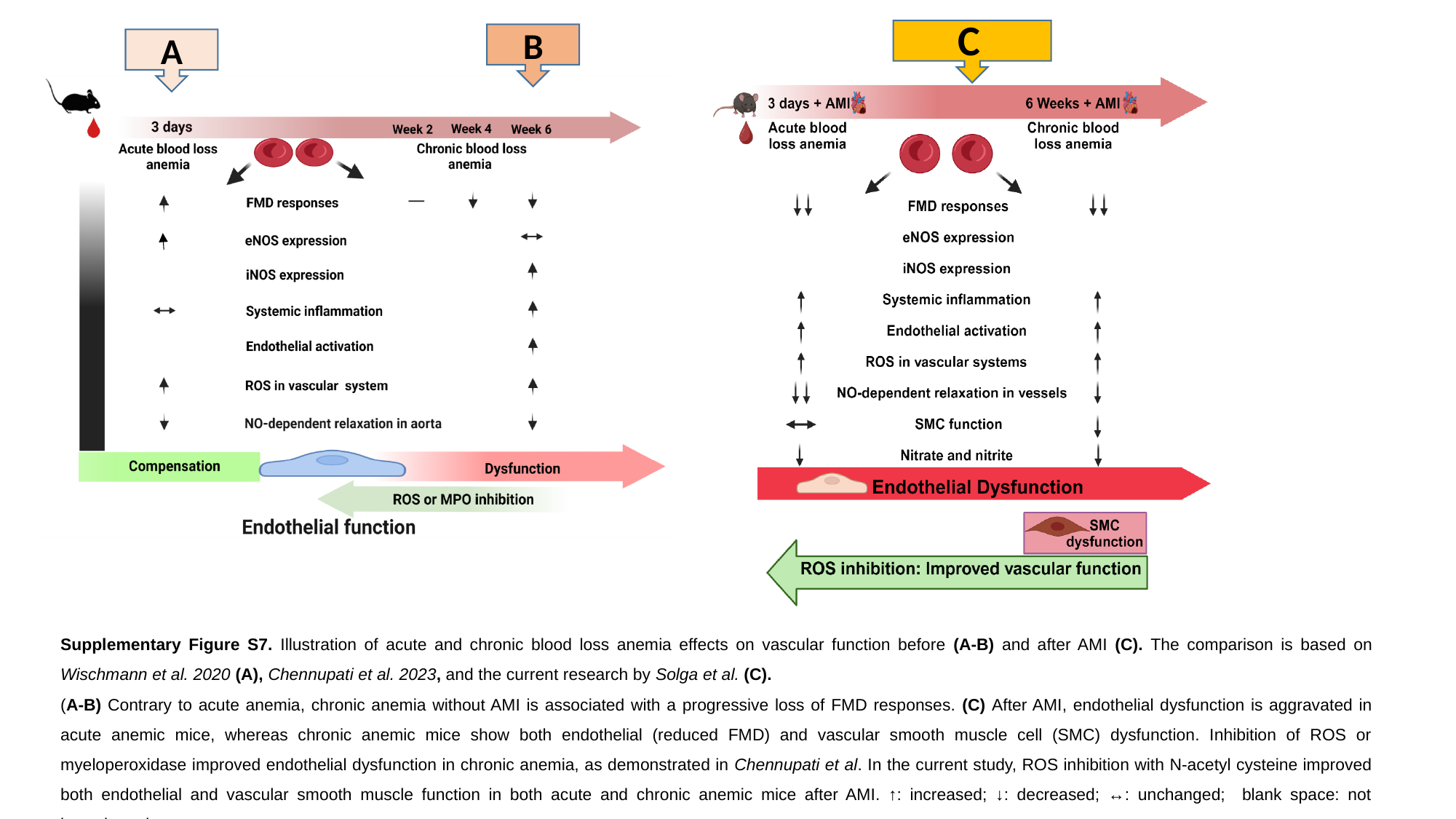

C
A
B
B
A
Supplementary Figure S7. Illustration of acute and chronic blood loss anemia effects on vascular function before (A-B) and after AMI (C). The comparison is based on Wischmann et al. 2020 (A), Chennupati et al. 2023, and the current research by Solga et al. (C).
(A-B) Contrary to acute anemia, chronic anemia without AMI is associated with a progressive loss of FMD responses. (C) After AMI, endothelial dysfunction is aggravated in acute anemic mice, whereas chronic anemic mice show both endothelial (reduced FMD) and vascular smooth muscle cell (SMC) dysfunction. Inhibition of ROS or myeloperoxidase improved endothelial dysfunction in chronic anemia, as demonstrated in Chennupati et al. In the current study, ROS inhibition with N-acetyl cysteine improved both endothelial and vascular smooth muscle function in both acute and chronic anemic mice after AMI. ↑: increased; ↓: decreased; ↔: unchanged; blank space: not investigated.
