## Supplementary tables 1-6 for "Distinct effects of acute and chronic blood loss anemia on vascular function after acute myocardial infarction"

### Slide 1
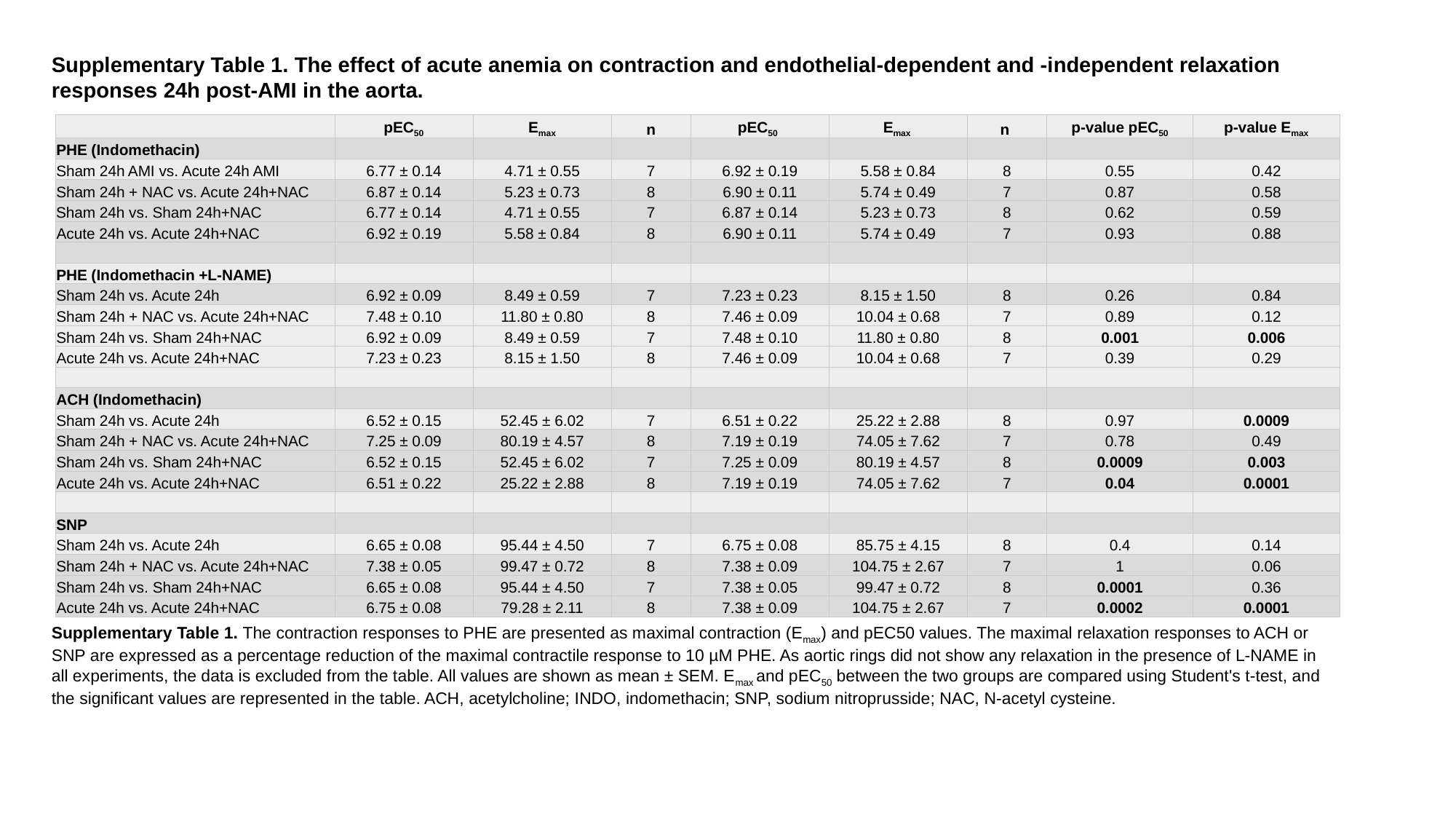

Supplementary Table 1. The effect of acute anemia on contraction and endothelial-dependent and -independent relaxation responses 24h post-AMI in the aorta.
| | pEC_50_ | E_max_ | n | pEC_50_ | E_max_ | n | p-value pEC_50_ | p-value E_max_ |
| --- | --- | --- | --- | --- | --- | --- | --- | --- |
| PHE (Indomethacin) | | | | | | | | |
| Sham 24h AMI vs. Acute 24h AMI | 6.77 ± 0.14 | 4.71 ± 0.55 | 7 | 6.92 ± 0.19 | 5.58 ± 0.84 | 8 | 0.55 | 0.42 |
| Sham 24h + NAC vs. Acute 24h+NAC | 6.87 ± 0.14 | 5.23 ± 0.73 | 8 | 6.90 ± 0.11 | 5.74 ± 0.49 | 7 | 0.87 | 0.58 |
| Sham 24h vs. Sham 24h+NAC | 6.77 ± 0.14 | 4.71 ± 0.55 | 7 | 6.87 ± 0.14 | 5.23 ± 0.73 | 8 | 0.62 | 0.59 |
| Acute 24h vs. Acute 24h+NAC | 6.92 ± 0.19 | 5.58 ± 0.84 | 8 | 6.90 ± 0.11 | 5.74 ± 0.49 | 7 | 0.93 | 0.88 |
| PHE (Indomethacin +L-NAME) | | | | | | | | |
| Sham 24h vs. Acute 24h | 6.92 ± 0.09 | 8.49 ± 0.59 | 7 | 7.23 ± 0.23 | 8.15 ± 1.50 | 8 | 0.26 | 0.84 |
| Sham 24h + NAC vs. Acute 24h+NAC | 7.48 ± 0.10 | 11.80 ± 0.80 | 8 | 7.46 ± 0.09 | 10.04 ± 0.68 | 7 | 0.89 | 0.12 |
| Sham 24h vs. Sham 24h+NAC | 6.92 ± 0.09 | 8.49 ± 0.59 | 7 | 7.48 ± 0.10 | 11.80 ± 0.80 | 8 | 0.001 | 0.006 |
| Acute 24h vs. Acute 24h+NAC | 7.23 ± 0.23 | 8.15 ± 1.50 | 8 | 7.46 ± 0.09 | 10.04 ± 0.68 | 7 | 0.39 | 0.29 |
| ACH (Indomethacin) | | | | | | | | |
| Sham 24h vs. Acute 24h | 6.52 ± 0.15 | 52.45 ± 6.02 | 7 | 6.51 ± 0.22 | 25.22 ± 2.88 | 8 | 0.97 | 0.0009 |
| Sham 24h + NAC vs. Acute 24h+NAC | 7.25 ± 0.09 | 80.19 ± 4.57 | 8 | 7.19 ± 0.19 | 74.05 ± 7.62 | 7 | 0.78 | 0.49 |
| Sham 24h vs. Sham 24h+NAC | 6.52 ± 0.15 | 52.45 ± 6.02 | 7 | 7.25 ± 0.09 | 80.19 ± 4.57 | 8 | 0.0009 | 0.003 |
| Acute 24h vs. Acute 24h+NAC | 6.51 ± 0.22 | 25.22 ± 2.88 | 8 | 7.19 ± 0.19 | 74.05 ± 7.62 | 7 | 0.04 | 0.0001 |
| SNP | | | | | | | | |
| Sham 24h vs. Acute 24h | 6.65 ± 0.08 | 95.44 ± 4.50 | 7 | 6.75 ± 0.08 | 85.75 ± 4.15 | 8 | 0.4 | 0.14 |
| Sham 24h + NAC vs. Acute 24h+NAC | 7.38 ± 0.05 | 99.47 ± 0.72 | 8 | 7.38 ± 0.09 | 104.75 ± 2.67 | 7 | 1 | 0.06 |
| Sham 24h vs. Sham 24h+NAC | 6.65 ± 0.08 | 95.44 ± 4.50 | 7 | 7.38 ± 0.05 | 99.47 ± 0.72 | 8 | 0.0001 | 0.36 |
| Acute 24h vs. Acute 24h+NAC | 6.75 ± 0.08 | 79.28 ± 2.11 | 8 | 7.38 ± 0.09 | 104.75 ± 2.67 | 7 | 0.0002 | 0.0001 |
Supplementary Table 1. The contraction responses to PHE are presented as maximal contraction (E_max_) and pEC50 values. The maximal relaxation responses to ACH or SNP are expressed as a percentage reduction of the maximal contractile response to 10 µM PHE. As aortic rings did not show any relaxation in the presence of L-NAME in all experiments, the data is excluded from the table. All values are shown as mean ± SEM. E_max_ and pEC_50_ between the two groups are compared using Student's t-test, and the significant values are represented in the table. ACH, acetylcholine; INDO, indomethacin; SNP, sodium nitroprusside; NAC, N-acetyl cysteine.

### Slide 2
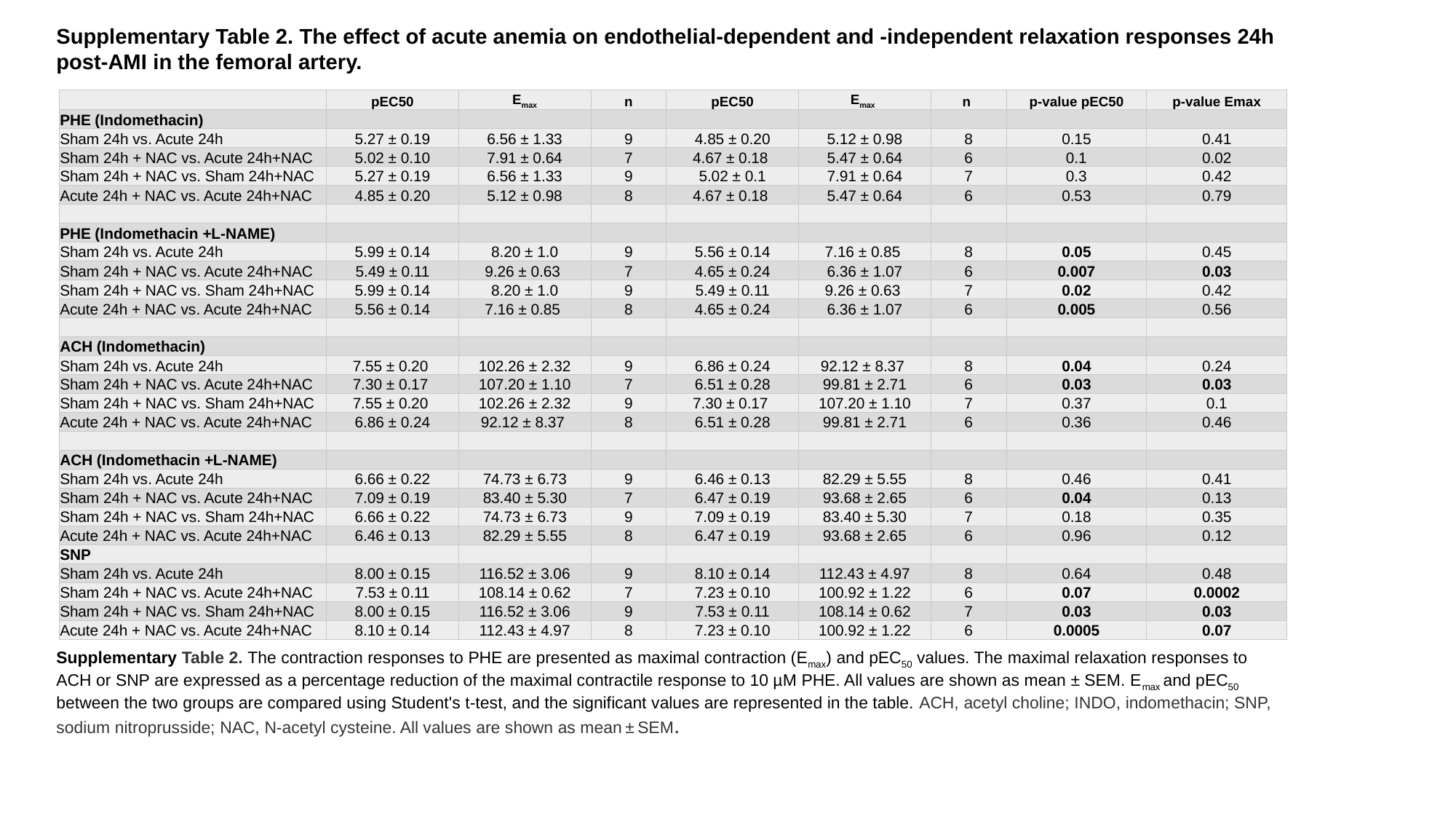

Supplementary Table 2. The effect of acute anemia on endothelial-dependent and -independent relaxation responses 24h post-AMI in the femoral artery.
| | pEC50 | E_max_ | n | pEC50 | E_max_ | n | p-value pEC50 | p-value Emax |
| --- | --- | --- | --- | --- | --- | --- | --- | --- |
| PHE (Indomethacin) | | | | | | | | |
| Sham 24h vs. Acute 24h | 5.27 ± 0.19 | 6.56 ± 1.33 | 9 | 4.85 ± 0.20 | 5.12 ± 0.98 | 8 | 0.15 | 0.41 |
| Sham 24h + NAC vs. Acute 24h+NAC | 5.02 ± 0.10 | 7.91 ± 0.64 | 7 | 4.67 ± 0.18 | 5.47 ± 0.64 | 6 | 0.1 | 0.02 |
| Sham 24h + NAC vs. Sham 24h+NAC | 5.27 ± 0.19 | 6.56 ± 1.33 | 9 | 5.02 ± 0.1 | 7.91 ± 0.64 | 7 | 0.3 | 0.42 |
| Acute 24h + NAC vs. Acute 24h+NAC | 4.85 ± 0.20 | 5.12 ± 0.98 | 8 | 4.67 ± 0.18 | 5.47 ± 0.64 | 6 | 0.53 | 0.79 |
| PHE (Indomethacin +L-NAME) | | | | | | | | |
| Sham 24h vs. Acute 24h | 5.99 ± 0.14 | 8.20 ± 1.0 | 9 | 5.56 ± 0.14 | 7.16 ± 0.85 | 8 | 0.05 | 0.45 |
| Sham 24h + NAC vs. Acute 24h+NAC | 5.49 ± 0.11 | 9.26 ± 0.63 | 7 | 4.65 ± 0.24 | 6.36 ± 1.07 | 6 | 0.007 | 0.03 |
| Sham 24h + NAC vs. Sham 24h+NAC | 5.99 ± 0.14 | 8.20 ± 1.0 | 9 | 5.49 ± 0.11 | 9.26 ± 0.63 | 7 | 0.02 | 0.42 |
| Acute 24h + NAC vs. Acute 24h+NAC | 5.56 ± 0.14 | 7.16 ± 0.85 | 8 | 4.65 ± 0.24 | 6.36 ± 1.07 | 6 | 0.005 | 0.56 |
| ACH (Indomethacin) | | | | | | | | |
| Sham 24h vs. Acute 24h | 7.55 ± 0.20 | 102.26 ± 2.32 | 9 | 6.86 ± 0.24 | 92.12 ± 8.37 | 8 | 0.04 | 0.24 |
| Sham 24h + NAC vs. Acute 24h+NAC | 7.30 ± 0.17 | 107.20 ± 1.10 | 7 | 6.51 ± 0.28 | 99.81 ± 2.71 | 6 | 0.03 | 0.03 |
| Sham 24h + NAC vs. Sham 24h+NAC | 7.55 ± 0.20 | 102.26 ± 2.32 | 9 | 7.30 ± 0.17 | 107.20 ± 1.10 | 7 | 0.37 | 0.1 |
| Acute 24h + NAC vs. Acute 24h+NAC | 6.86 ± 0.24 | 92.12 ± 8.37 | 8 | 6.51 ± 0.28 | 99.81 ± 2.71 | 6 | 0.36 | 0.46 |
| ACH (Indomethacin +L-NAME) | | | | | | | | |
| Sham 24h vs. Acute 24h | 6.66 ± 0.22 | 74.73 ± 6.73 | 9 | 6.46 ± 0.13 | 82.29 ± 5.55 | 8 | 0.46 | 0.41 |
| Sham 24h + NAC vs. Acute 24h+NAC | 7.09 ± 0.19 | 83.40 ± 5.30 | 7 | 6.47 ± 0.19 | 93.68 ± 2.65 | 6 | 0.04 | 0.13 |
| Sham 24h + NAC vs. Sham 24h+NAC | 6.66 ± 0.22 | 74.73 ± 6.73 | 9 | 7.09 ± 0.19 | 83.40 ± 5.30 | 7 | 0.18 | 0.35 |
| Acute 24h + NAC vs. Acute 24h+NAC | 6.46 ± 0.13 | 82.29 ± 5.55 | 8 | 6.47 ± 0.19 | 93.68 ± 2.65 | 6 | 0.96 | 0.12 |
| SNP | | | | | | | | |
| Sham 24h vs. Acute 24h | 8.00 ± 0.15 | 116.52 ± 3.06 | 9 | 8.10 ± 0.14 | 112.43 ± 4.97 | 8 | 0.64 | 0.48 |
| Sham 24h + NAC vs. Acute 24h+NAC | 7.53 ± 0.11 | 108.14 ± 0.62 | 7 | 7.23 ± 0.10 | 100.92 ± 1.22 | 6 | 0.07 | 0.0002 |
| Sham 24h + NAC vs. Sham 24h+NAC | 8.00 ± 0.15 | 116.52 ± 3.06 | 9 | 7.53 ± 0.11 | 108.14 ± 0.62 | 7 | 0.03 | 0.03 |
| Acute 24h + NAC vs. Acute 24h+NAC | 8.10 ± 0.14 | 112.43 ± 4.97 | 8 | 7.23 ± 0.10 | 100.92 ± 1.22 | 6 | 0.0005 | 0.07 |
Supplementary Table 2. The contraction responses to PHE are presented as maximal contraction (E_max_) and pEC_50_ values. The maximal relaxation responses to ACH or SNP are expressed as a percentage reduction of the maximal contractile response to 10 µM PHE. All values are shown as mean ± SEM. E_max_ and pEC_50_ between the two groups are compared using Student's t-test, and the significant values are represented in the table. ACH, acetyl choline; INDO, indomethacin; SNP, sodium nitroprusside; NAC, N-acetyl cysteine. All values are shown as mean ± SEM.

### Slide 3
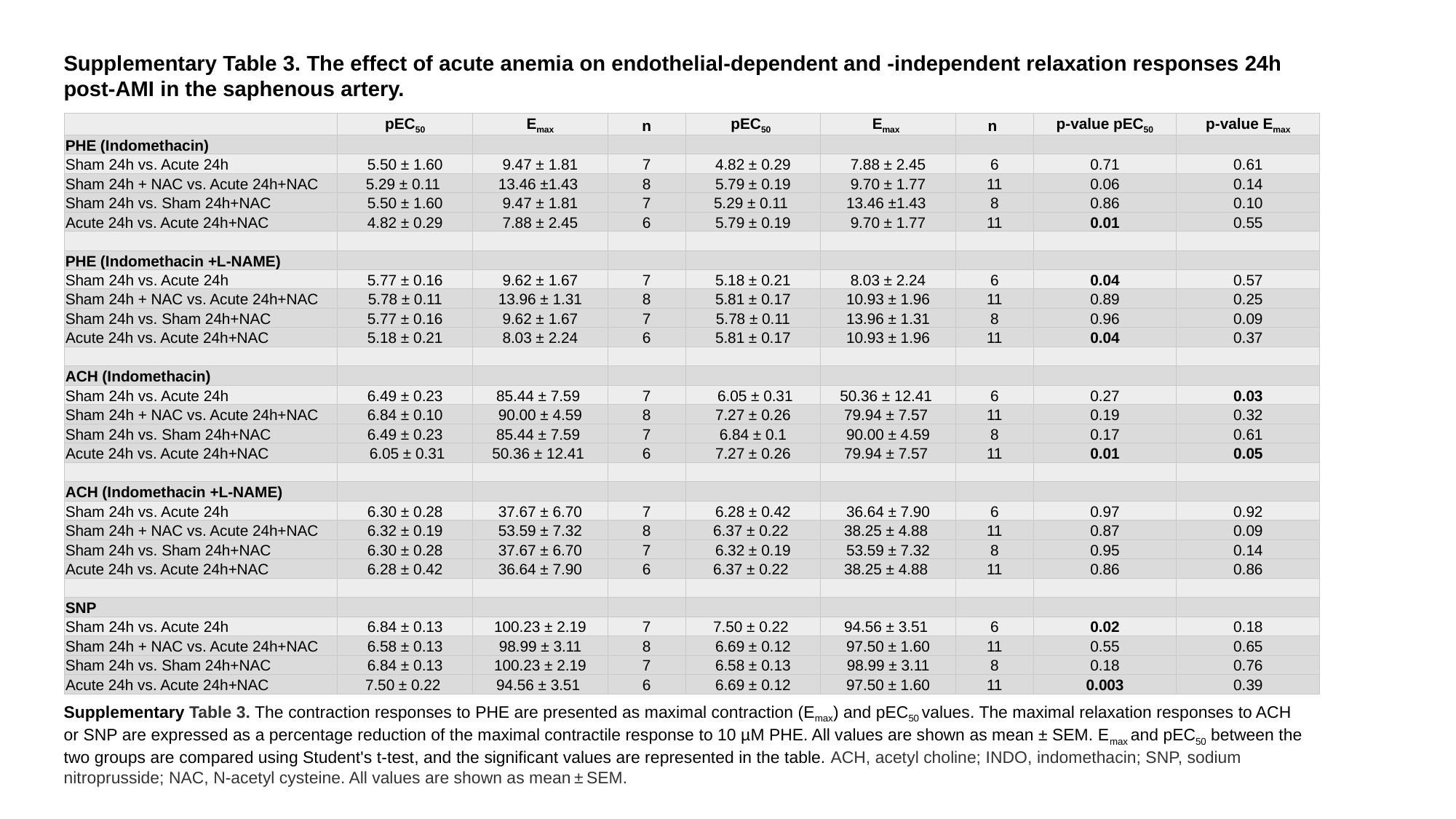

Supplementary Table 3. The effect of acute anemia on endothelial-dependent and -independent relaxation responses 24h post-AMI in the saphenous artery.
| | pEC_50_ | E_max_ | n | pEC_50_ | E_max_ | n | p-value pEC_50_ | p-value E_max_ |
| --- | --- | --- | --- | --- | --- | --- | --- | --- |
| PHE (Indomethacin) | | | | | | | | |
| Sham 24h vs. Acute 24h | 5.50 ± 1.60 | 9.47 ± 1.81 | 7 | 4.82 ± 0.29 | 7.88 ± 2.45 | 6 | 0.71 | 0.61 |
| Sham 24h + NAC vs. Acute 24h+NAC | 5.29 ± 0.11 | 13.46 ±1.43 | 8 | 5.79 ± 0.19 | 9.70 ± 1.77 | 11 | 0.06 | 0.14 |
| Sham 24h vs. Sham 24h+NAC | 5.50 ± 1.60 | 9.47 ± 1.81 | 7 | 5.29 ± 0.11 | 13.46 ±1.43 | 8 | 0.86 | 0.10 |
| Acute 24h vs. Acute 24h+NAC | 4.82 ± 0.29 | 7.88 ± 2.45 | 6 | 5.79 ± 0.19 | 9.70 ± 1.77 | 11 | 0.01 | 0.55 |
| PHE (Indomethacin +L-NAME) | | | | | | | | |
| Sham 24h vs. Acute 24h | 5.77 ± 0.16 | 9.62 ± 1.67 | 7 | 5.18 ± 0.21 | 8.03 ± 2.24 | 6 | 0.04 | 0.57 |
| Sham 24h + NAC vs. Acute 24h+NAC | 5.78 ± 0.11 | 13.96 ± 1.31 | 8 | 5.81 ± 0.17 | 10.93 ± 1.96 | 11 | 0.89 | 0.25 |
| Sham 24h vs. Sham 24h+NAC | 5.77 ± 0.16 | 9.62 ± 1.67 | 7 | 5.78 ± 0.11 | 13.96 ± 1.31 | 8 | 0.96 | 0.09 |
| Acute 24h vs. Acute 24h+NAC | 5.18 ± 0.21 | 8.03 ± 2.24 | 6 | 5.81 ± 0.17 | 10.93 ± 1.96 | 11 | 0.04 | 0.37 |
| ACH (Indomethacin) | | | | | | | | |
| Sham 24h vs. Acute 24h | 6.49 ± 0.23 | 85.44 ± 7.59 | 7 | 6.05 ± 0.31 | 50.36 ± 12.41 | 6 | 0.27 | 0.03 |
| Sham 24h + NAC vs. Acute 24h+NAC | 6.84 ± 0.10 | 90.00 ± 4.59 | 8 | 7.27 ± 0.26 | 79.94 ± 7.57 | 11 | 0.19 | 0.32 |
| Sham 24h vs. Sham 24h+NAC | 6.49 ± 0.23 | 85.44 ± 7.59 | 7 | 6.84 ± 0.1 | 90.00 ± 4.59 | 8 | 0.17 | 0.61 |
| Acute 24h vs. Acute 24h+NAC | 6.05 ± 0.31 | 50.36 ± 12.41 | 6 | 7.27 ± 0.26 | 79.94 ± 7.57 | 11 | 0.01 | 0.05 |
| ACH (Indomethacin +L-NAME) | | | | | | | | |
| Sham 24h vs. Acute 24h | 6.30 ± 0.28 | 37.67 ± 6.70 | 7 | 6.28 ± 0.42 | 36.64 ± 7.90 | 6 | 0.97 | 0.92 |
| Sham 24h + NAC vs. Acute 24h+NAC | 6.32 ± 0.19 | 53.59 ± 7.32 | 8 | 6.37 ± 0.22 | 38.25 ± 4.88 | 11 | 0.87 | 0.09 |
| Sham 24h vs. Sham 24h+NAC | 6.30 ± 0.28 | 37.67 ± 6.70 | 7 | 6.32 ± 0.19 | 53.59 ± 7.32 | 8 | 0.95 | 0.14 |
| Acute 24h vs. Acute 24h+NAC | 6.28 ± 0.42 | 36.64 ± 7.90 | 6 | 6.37 ± 0.22 | 38.25 ± 4.88 | 11 | 0.86 | 0.86 |
| SNP | | | | | | | | |
| Sham 24h vs. Acute 24h | 6.84 ± 0.13 | 100.23 ± 2.19 | 7 | 7.50 ± 0.22 | 94.56 ± 3.51 | 6 | 0.02 | 0.18 |
| Sham 24h + NAC vs. Acute 24h+NAC | 6.58 ± 0.13 | 98.99 ± 3.11 | 8 | 6.69 ± 0.12 | 97.50 ± 1.60 | 11 | 0.55 | 0.65 |
| Sham 24h vs. Sham 24h+NAC | 6.84 ± 0.13 | 100.23 ± 2.19 | 7 | 6.58 ± 0.13 | 98.99 ± 3.11 | 8 | 0.18 | 0.76 |
| Acute 24h vs. Acute 24h+NAC | 7.50 ± 0.22 | 94.56 ± 3.51 | 6 | 6.69 ± 0.12 | 97.50 ± 1.60 | 11 | 0.003 | 0.39 |
Supplementary Table 3. The contraction responses to PHE are presented as maximal contraction (E_max_) and pEC_50_ values. The maximal relaxation responses to ACH or SNP are expressed as a percentage reduction of the maximal contractile response to 10 µM PHE. All values are shown as mean ± SEM. E_max_ and pEC_50_ between the two groups are compared using Student's t-test, and the significant values are represented in the table. ACH, acetyl choline; INDO, indomethacin; SNP, sodium nitroprusside; NAC, N-acetyl cysteine. All values are shown as mean ± SEM.

### Slide 4
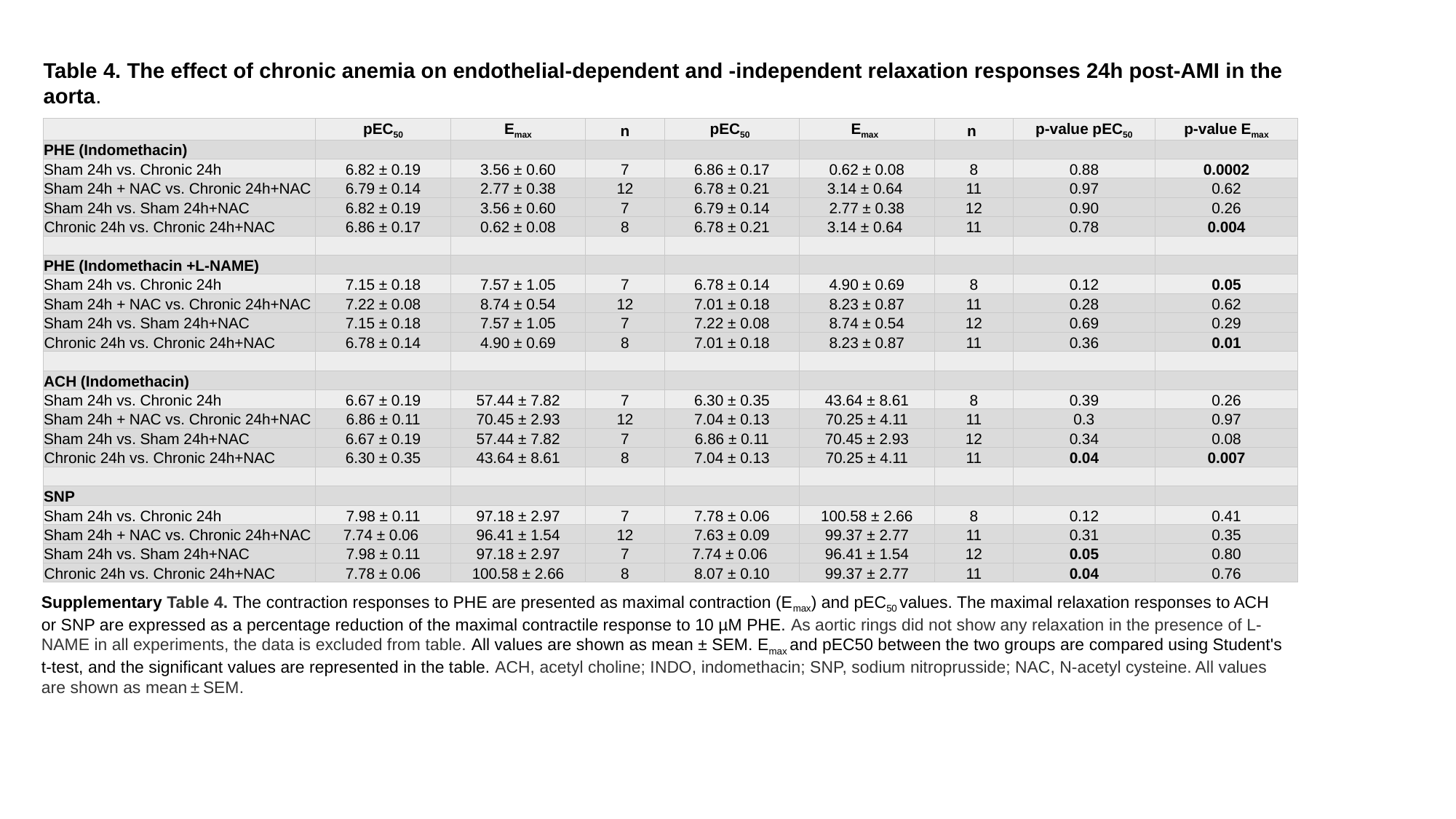

Table 4. The effect of chronic anemia on endothelial-dependent and -independent relaxation responses 24h post-AMI in the aorta.
| | pEC_50_ | E_max_ | n | pEC_50_ | E_max_ | n | p-value pEC_50_ | p-value E_max_ |
| --- | --- | --- | --- | --- | --- | --- | --- | --- |
| PHE (Indomethacin) | | | | | | | | |
| Sham 24h vs. Chronic 24h | 6.82 ± 0.19 | 3.56 ± 0.60 | 7 | 6.86 ± 0.17 | 0.62 ± 0.08 | 8 | 0.88 | 0.0002 |
| Sham 24h + NAC vs. Chronic 24h+NAC | 6.79 ± 0.14 | 2.77 ± 0.38 | 12 | 6.78 ± 0.21 | 3.14 ± 0.64 | 11 | 0.97 | 0.62 |
| Sham 24h vs. Sham 24h+NAC | 6.82 ± 0.19 | 3.56 ± 0.60 | 7 | 6.79 ± 0.14 | 2.77 ± 0.38 | 12 | 0.90 | 0.26 |
| Chronic 24h vs. Chronic 24h+NAC | 6.86 ± 0.17 | 0.62 ± 0.08 | 8 | 6.78 ± 0.21 | 3.14 ± 0.64 | 11 | 0.78 | 0.004 |
| PHE (Indomethacin +L-NAME) | | | | | | | | |
| Sham 24h vs. Chronic 24h | 7.15 ± 0.18 | 7.57 ± 1.05 | 7 | 6.78 ± 0.14 | 4.90 ± 0.69 | 8 | 0.12 | 0.05 |
| Sham 24h + NAC vs. Chronic 24h+NAC | 7.22 ± 0.08 | 8.74 ± 0.54 | 12 | 7.01 ± 0.18 | 8.23 ± 0.87 | 11 | 0.28 | 0.62 |
| Sham 24h vs. Sham 24h+NAC | 7.15 ± 0.18 | 7.57 ± 1.05 | 7 | 7.22 ± 0.08 | 8.74 ± 0.54 | 12 | 0.69 | 0.29 |
| Chronic 24h vs. Chronic 24h+NAC | 6.78 ± 0.14 | 4.90 ± 0.69 | 8 | 7.01 ± 0.18 | 8.23 ± 0.87 | 11 | 0.36 | 0.01 |
| ACH (Indomethacin) | | | | | | | | |
| Sham 24h vs. Chronic 24h | 6.67 ± 0.19 | 57.44 ± 7.82 | 7 | 6.30 ± 0.35 | 43.64 ± 8.61 | 8 | 0.39 | 0.26 |
| Sham 24h + NAC vs. Chronic 24h+NAC | 6.86 ± 0.11 | 70.45 ± 2.93 | 12 | 7.04 ± 0.13 | 70.25 ± 4.11 | 11 | 0.3 | 0.97 |
| Sham 24h vs. Sham 24h+NAC | 6.67 ± 0.19 | 57.44 ± 7.82 | 7 | 6.86 ± 0.11 | 70.45 ± 2.93 | 12 | 0.34 | 0.08 |
| Chronic 24h vs. Chronic 24h+NAC | 6.30 ± 0.35 | 43.64 ± 8.61 | 8 | 7.04 ± 0.13 | 70.25 ± 4.11 | 11 | 0.04 | 0.007 |
| SNP | | | | | | | | |
| Sham 24h vs. Chronic 24h | 7.98 ± 0.11 | 97.18 ± 2.97 | 7 | 7.78 ± 0.06 | 100.58 ± 2.66 | 8 | 0.12 | 0.41 |
| Sham 24h + NAC vs. Chronic 24h+NAC | 7.74 ± 0.06 | 96.41 ± 1.54 | 12 | 7.63 ± 0.09 | 99.37 ± 2.77 | 11 | 0.31 | 0.35 |
| Sham 24h vs. Sham 24h+NAC | 7.98 ± 0.11 | 97.18 ± 2.97 | 7 | 7.74 ± 0.06 | 96.41 ± 1.54 | 12 | 0.05 | 0.80 |
| Chronic 24h vs. Chronic 24h+NAC | 7.78 ± 0.06 | 100.58 ± 2.66 | 8 | 8.07 ± 0.10 | 99.37 ± 2.77 | 11 | 0.04 | 0.76 |
Supplementary Table 4. The contraction responses to PHE are presented as maximal contraction (E_max_) and pEC_50_ values. The maximal relaxation responses to ACH or SNP are expressed as a percentage reduction of the maximal contractile response to 10 µM PHE. As aortic rings did not show any relaxation in the presence of L-NAME in all experiments, the data is excluded from table. All values are shown as mean ± SEM. E_max_ and pEC50 between the two groups are compared using Student's t-test, and the significant values are represented in the table. ACH, acetyl choline; INDO, indomethacin; SNP, sodium nitroprusside; NAC, N-acetyl cysteine. All values are shown as mean ± SEM.

### Slide 5
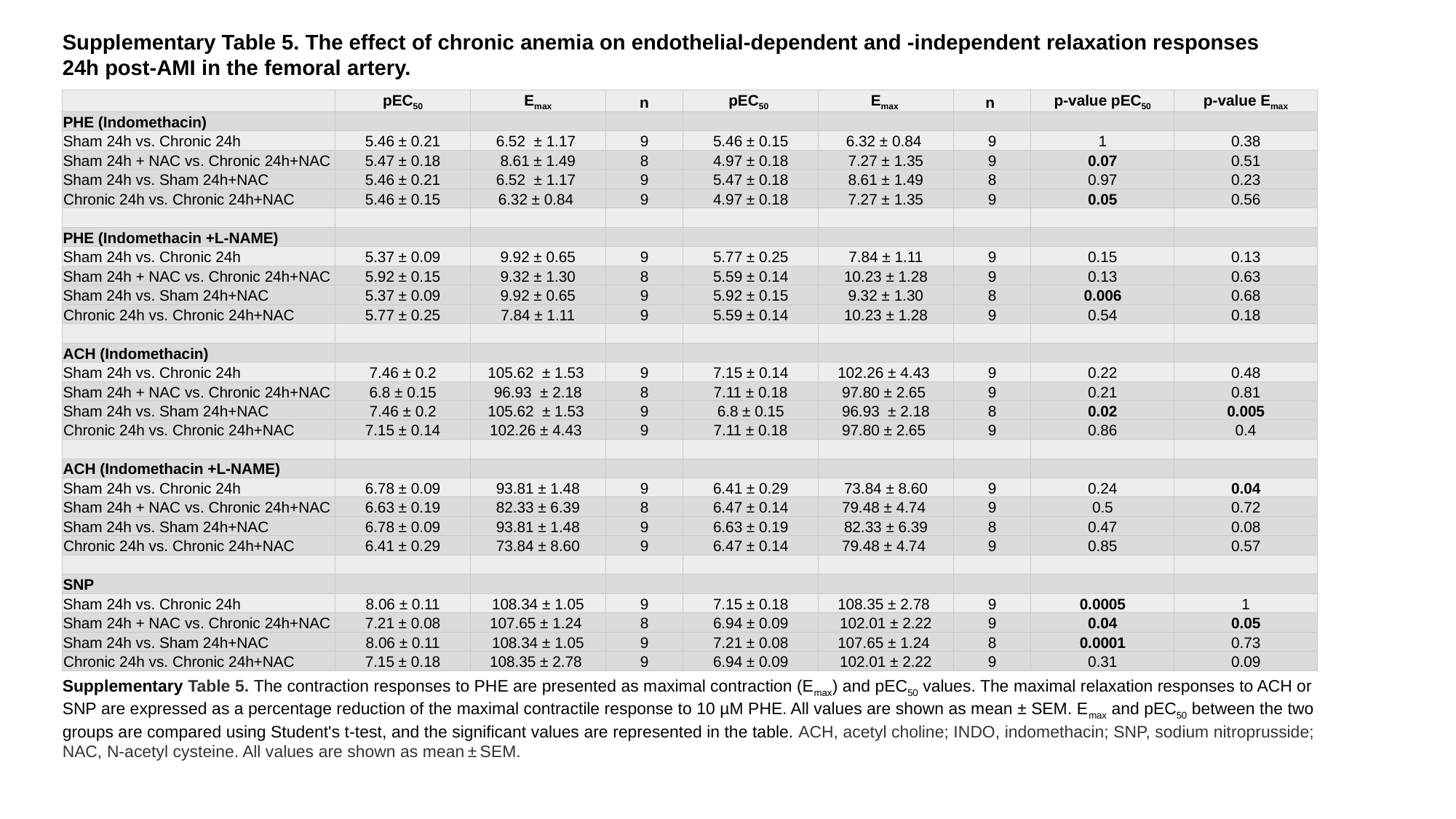

Supplementary Table 5. The effect of chronic anemia on endothelial-dependent and -independent relaxation responses 24h post-AMI in the femoral artery.
| | pEC_50_ | E_max_ | n | pEC_50_ | E_max_ | n | p-value pEC_50_ | p-value E_max_ |
| --- | --- | --- | --- | --- | --- | --- | --- | --- |
| PHE (Indomethacin) | | | | | | | | |
| Sham 24h vs. Chronic 24h | 5.46 ± 0.21 | 6.52 ± 1.17 | 9 | 5.46 ± 0.15 | 6.32 ± 0.84 | 9 | 1 | 0.38 |
| Sham 24h + NAC vs. Chronic 24h+NAC | 5.47 ± 0.18 | 8.61 ± 1.49 | 8 | 4.97 ± 0.18 | 7.27 ± 1.35 | 9 | 0.07 | 0.51 |
| Sham 24h vs. Sham 24h+NAC | 5.46 ± 0.21 | 6.52 ± 1.17 | 9 | 5.47 ± 0.18 | 8.61 ± 1.49 | 8 | 0.97 | 0.23 |
| Chronic 24h vs. Chronic 24h+NAC | 5.46 ± 0.15 | 6.32 ± 0.84 | 9 | 4.97 ± 0.18 | 7.27 ± 1.35 | 9 | 0.05 | 0.56 |
| PHE (Indomethacin +L-NAME) | | | | | | | | |
| Sham 24h vs. Chronic 24h | 5.37 ± 0.09 | 9.92 ± 0.65 | 9 | 5.77 ± 0.25 | 7.84 ± 1.11 | 9 | 0.15 | 0.13 |
| Sham 24h + NAC vs. Chronic 24h+NAC | 5.92 ± 0.15 | 9.32 ± 1.30 | 8 | 5.59 ± 0.14 | 10.23 ± 1.28 | 9 | 0.13 | 0.63 |
| Sham 24h vs. Sham 24h+NAC | 5.37 ± 0.09 | 9.92 ± 0.65 | 9 | 5.92 ± 0.15 | 9.32 ± 1.30 | 8 | 0.006 | 0.68 |
| Chronic 24h vs. Chronic 24h+NAC | 5.77 ± 0.25 | 7.84 ± 1.11 | 9 | 5.59 ± 0.14 | 10.23 ± 1.28 | 9 | 0.54 | 0.18 |
| ACH (Indomethacin) | | | | | | | | |
| Sham 24h vs. Chronic 24h | 7.46 ± 0.2 | 105.62 ± 1.53 | 9 | 7.15 ± 0.14 | 102.26 ± 4.43 | 9 | 0.22 | 0.48 |
| Sham 24h + NAC vs. Chronic 24h+NAC | 6.8 ± 0.15 | 96.93 ± 2.18 | 8 | 7.11 ± 0.18 | 97.80 ± 2.65 | 9 | 0.21 | 0.81 |
| Sham 24h vs. Sham 24h+NAC | 7.46 ± 0.2 | 105.62 ± 1.53 | 9 | 6.8 ± 0.15 | 96.93 ± 2.18 | 8 | 0.02 | 0.005 |
| Chronic 24h vs. Chronic 24h+NAC | 7.15 ± 0.14 | 102.26 ± 4.43 | 9 | 7.11 ± 0.18 | 97.80 ± 2.65 | 9 | 0.86 | 0.4 |
| ACH (Indomethacin +L-NAME) | | | | | | | | |
| Sham 24h vs. Chronic 24h | 6.78 ± 0.09 | 93.81 ± 1.48 | 9 | 6.41 ± 0.29 | 73.84 ± 8.60 | 9 | 0.24 | 0.04 |
| Sham 24h + NAC vs. Chronic 24h+NAC | 6.63 ± 0.19 | 82.33 ± 6.39 | 8 | 6.47 ± 0.14 | 79.48 ± 4.74 | 9 | 0.5 | 0.72 |
| Sham 24h vs. Sham 24h+NAC | 6.78 ± 0.09 | 93.81 ± 1.48 | 9 | 6.63 ± 0.19 | 82.33 ± 6.39 | 8 | 0.47 | 0.08 |
| Chronic 24h vs. Chronic 24h+NAC | 6.41 ± 0.29 | 73.84 ± 8.60 | 9 | 6.47 ± 0.14 | 79.48 ± 4.74 | 9 | 0.85 | 0.57 |
| SNP | | | | | | | | |
| Sham 24h vs. Chronic 24h | 8.06 ± 0.11 | 108.34 ± 1.05 | 9 | 7.15 ± 0.18 | 108.35 ± 2.78 | 9 | 0.0005 | 1 |
| Sham 24h + NAC vs. Chronic 24h+NAC | 7.21 ± 0.08 | 107.65 ± 1.24 | 8 | 6.94 ± 0.09 | 102.01 ± 2.22 | 9 | 0.04 | 0.05 |
| Sham 24h vs. Sham 24h+NAC | 8.06 ± 0.11 | 108.34 ± 1.05 | 9 | 7.21 ± 0.08 | 107.65 ± 1.24 | 8 | 0.0001 | 0.73 |
| Chronic 24h vs. Chronic 24h+NAC | 7.15 ± 0.18 | 108.35 ± 2.78 | 9 | 6.94 ± 0.09 | 102.01 ± 2.22 | 9 | 0.31 | 0.09 |
Supplementary Table 5. The contraction responses to PHE are presented as maximal contraction (E_max_) and pEC_50_ values. The maximal relaxation responses to ACH or SNP are expressed as a percentage reduction of the maximal contractile response to 10 µM PHE. All values are shown as mean ± SEM. E_max_ and pEC_50_ between the two groups are compared using Student's t-test, and the significant values are represented in the table. ACH, acetyl choline; INDO, indomethacin; SNP, sodium nitroprusside; NAC, N-acetyl cysteine. All values are shown as mean ± SEM.

### Slide 6
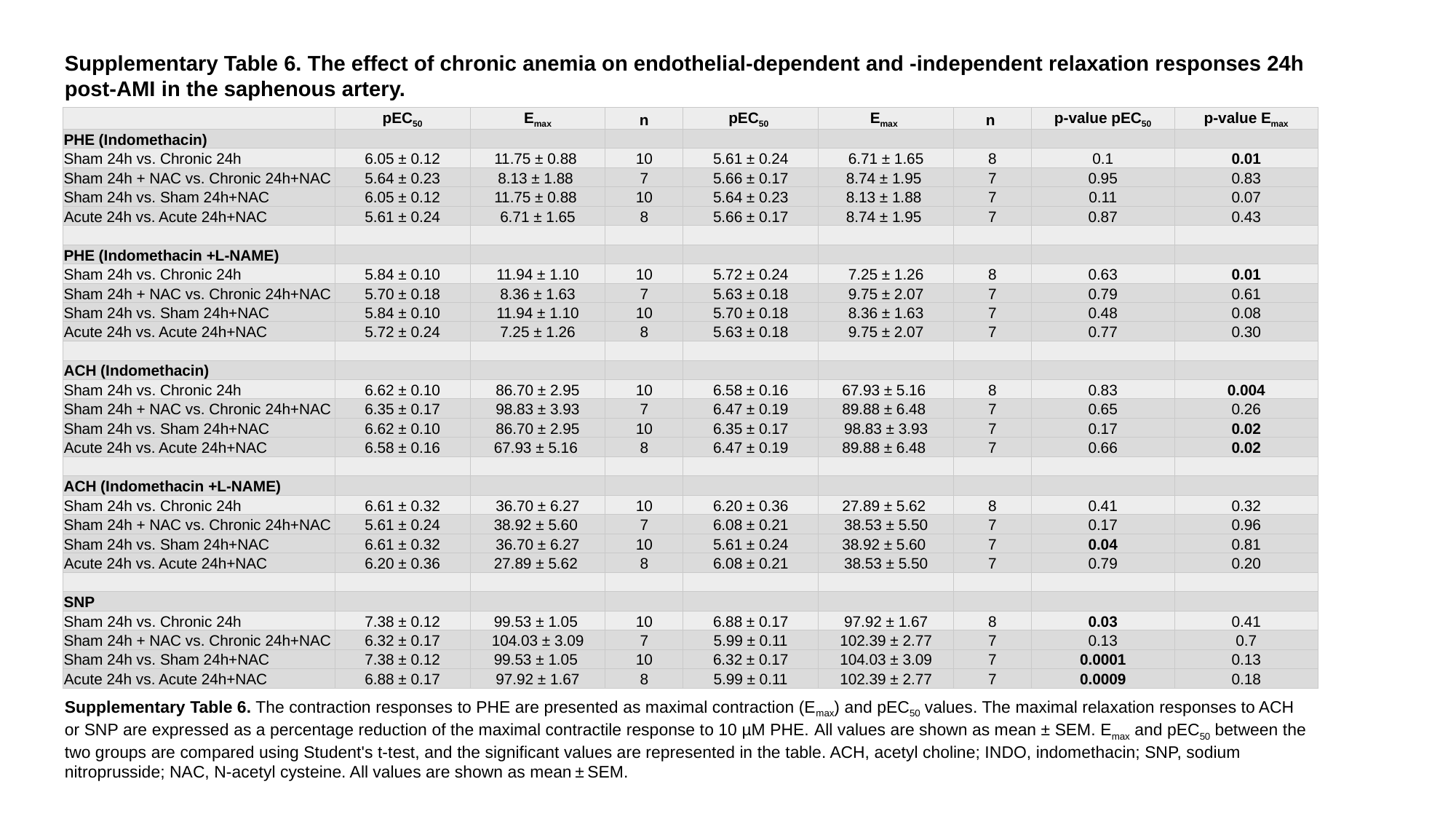

Supplementary Table 6. The effect of chronic anemia on endothelial-dependent and -independent relaxation responses 24h post-AMI in the saphenous artery.
| | pEC_50_ | E_max_ | n | pEC_50_ | E_max_ | n | p-value pEC_50_ | p-value E_max_ |
| --- | --- | --- | --- | --- | --- | --- | --- | --- |
| PHE (Indomethacin) | | | | | | | | |
| Sham 24h vs. Chronic 24h | 6.05 ± 0.12 | 11.75 ± 0.88 | 10 | 5.61 ± 0.24 | 6.71 ± 1.65 | 8 | 0.1 | 0.01 |
| Sham 24h + NAC vs. Chronic 24h+NAC | 5.64 ± 0.23 | 8.13 ± 1.88 | 7 | 5.66 ± 0.17 | 8.74 ± 1.95 | 7 | 0.95 | 0.83 |
| Sham 24h vs. Sham 24h+NAC | 6.05 ± 0.12 | 11.75 ± 0.88 | 10 | 5.64 ± 0.23 | 8.13 ± 1.88 | 7 | 0.11 | 0.07 |
| Acute 24h vs. Acute 24h+NAC | 5.61 ± 0.24 | 6.71 ± 1.65 | 8 | 5.66 ± 0.17 | 8.74 ± 1.95 | 7 | 0.87 | 0.43 |
| PHE (Indomethacin +L-NAME) | | | | | | | | |
| Sham 24h vs. Chronic 24h | 5.84 ± 0.10 | 11.94 ± 1.10 | 10 | 5.72 ± 0.24 | 7.25 ± 1.26 | 8 | 0.63 | 0.01 |
| Sham 24h + NAC vs. Chronic 24h+NAC | 5.70 ± 0.18 | 8.36 ± 1.63 | 7 | 5.63 ± 0.18 | 9.75 ± 2.07 | 7 | 0.79 | 0.61 |
| Sham 24h vs. Sham 24h+NAC | 5.84 ± 0.10 | 11.94 ± 1.10 | 10 | 5.70 ± 0.18 | 8.36 ± 1.63 | 7 | 0.48 | 0.08 |
| Acute 24h vs. Acute 24h+NAC | 5.72 ± 0.24 | 7.25 ± 1.26 | 8 | 5.63 ± 0.18 | 9.75 ± 2.07 | 7 | 0.77 | 0.30 |
| ACH (Indomethacin) | | | | | | | | |
| Sham 24h vs. Chronic 24h | 6.62 ± 0.10 | 86.70 ± 2.95 | 10 | 6.58 ± 0.16 | 67.93 ± 5.16 | 8 | 0.83 | 0.004 |
| Sham 24h + NAC vs. Chronic 24h+NAC | 6.35 ± 0.17 | 98.83 ± 3.93 | 7 | 6.47 ± 0.19 | 89.88 ± 6.48 | 7 | 0.65 | 0.26 |
| Sham 24h vs. Sham 24h+NAC | 6.62 ± 0.10 | 86.70 ± 2.95 | 10 | 6.35 ± 0.17 | 98.83 ± 3.93 | 7 | 0.17 | 0.02 |
| Acute 24h vs. Acute 24h+NAC | 6.58 ± 0.16 | 67.93 ± 5.16 | 8 | 6.47 ± 0.19 | 89.88 ± 6.48 | 7 | 0.66 | 0.02 |
| ACH (Indomethacin +L-NAME) | | | | | | | | |
| Sham 24h vs. Chronic 24h | 6.61 ± 0.32 | 36.70 ± 6.27 | 10 | 6.20 ± 0.36 | 27.89 ± 5.62 | 8 | 0.41 | 0.32 |
| Sham 24h + NAC vs. Chronic 24h+NAC | 5.61 ± 0.24 | 38.92 ± 5.60 | 7 | 6.08 ± 0.21 | 38.53 ± 5.50 | 7 | 0.17 | 0.96 |
| Sham 24h vs. Sham 24h+NAC | 6.61 ± 0.32 | 36.70 ± 6.27 | 10 | 5.61 ± 0.24 | 38.92 ± 5.60 | 7 | 0.04 | 0.81 |
| Acute 24h vs. Acute 24h+NAC | 6.20 ± 0.36 | 27.89 ± 5.62 | 8 | 6.08 ± 0.21 | 38.53 ± 5.50 | 7 | 0.79 | 0.20 |
| SNP | | | | | | | | |
| Sham 24h vs. Chronic 24h | 7.38 ± 0.12 | 99.53 ± 1.05 | 10 | 6.88 ± 0.17 | 97.92 ± 1.67 | 8 | 0.03 | 0.41 |
| Sham 24h + NAC vs. Chronic 24h+NAC | 6.32 ± 0.17 | 104.03 ± 3.09 | 7 | 5.99 ± 0.11 | 102.39 ± 2.77 | 7 | 0.13 | 0.7 |
| Sham 24h vs. Sham 24h+NAC | 7.38 ± 0.12 | 99.53 ± 1.05 | 10 | 6.32 ± 0.17 | 104.03 ± 3.09 | 7 | 0.0001 | 0.13 |
| Acute 24h vs. Acute 24h+NAC | 6.88 ± 0.17 | 97.92 ± 1.67 | 8 | 5.99 ± 0.11 | 102.39 ± 2.77 | 7 | 0.0009 | 0.18 |
Supplementary Table 6. The contraction responses to PHE are presented as maximal contraction (E_max_) and pEC_50_ values. The maximal relaxation responses to ACH or SNP are expressed as a percentage reduction of the maximal contractile response to 10 µM PHE. All values are shown as mean ± SEM. E_max_ and pEC_50_ between the two groups are compared using Student's t-test, and the significant values are represented in the table. ACH, acetyl choline; INDO, indomethacin; SNP, sodium nitroprusside; NAC, N-acetyl cysteine. All values are shown as mean ± SEM.
